# Identification of the KIF18A alpha-4 helix as a therapeutic target for chromosomally unstable tumor cells

**DOI:** 10.1101/2023.10.16.562576

**Authors:** Katherine Schutt, Katelyn A. Queen, Kira Fisher, Olivia Budington, Weifeng Mao, Wei Liu, Yisong Xiao, Fred Aswad, James Joseph, Jason Stumpff

## Abstract

The mitotic kinesin, KIF18A, is required for proliferation of cancer cells that exhibit chromosome instability (CIN), implicating it as a promising target for treatment of a subset of aggressive tumor types. Determining regions of the KIF18A protein to target for inhibition will be important for the design and optimization of effective small molecule inhibitors. In this study, we investigated the effects of mutating S284 within the alpha-4 helix of KIF18A, which was previously identified as a phosphorylated residue. Mutations in S284 cause relocalization of KIF18A from the plus-ends of spindle microtubules to the spindle poles. Furthermore, KIF18A S284 mutants display loss of KIF18A function and fail to support proliferation in CIN tumor cells. Interestingly, similar effects on KIF18A localization and function were seen after treatment of CIN cells with KIF18A inhibitory compounds that are predicted to interact with residues within the alpha-4 helix. These data implicate the KIF18A alpha-4 helix as an effective target for inhibition and demonstrate that small molecules targeting KIF18A selectively limit CIN tumor cell proliferation and result in phenotypically similar effects on mitosis at the single cell level compared to genetic perturbations.

## INTRODUCTION

Faithful replication and segregation of chromosomes is essential for maintaining genomic integrity. A hallmark of tumor cells is chromosomal instability (CIN), where cells exhibit frequent gain or loss of whole chromosomes (1). These frequent missegregration events are in part due to inherent differences in CIN cells, such as increased microtubule plus-end assembly rates, aberrant kinetochore-microtubule attachments, and defects in the spindle assembly checkpoint (SAC) (2–4). Although CIN can lead to increased tumorigenesis and therapeutic resistance (5), recent efforts have focused on methods for targeting CIN as a possible Achilles’ heel to selectively target and eliminate tumor cells. Common therapeutic strategies for treatment of tumors directly target the altered microtubule dynamics of CIN cells with microtubule-targeting agents (MTAs) such as Paclitaxel, which stabilizes microtubules and induces mitotic arrest or aberrant chromosome segregation (6,7). Although many MTAs show promising results in pre-clinical studies, clinical application of MTAs in patients have caused off-target effects and cytotoxicity, necessitating additional therapeutic options (8,9).

Selectively targeting CIN requires identifying a genetic dependency of chromosomally unstable cells versus diploid cells. Several recent studies revealed a specific dependency of CIN cells, but not diploid cells, on the kinesin-8 KIF18A (10–12). KIF18A accumulates at the plus-ends of microtubules, where is suppresses microtubule dynamics to promote chromosome alignment during metaphase (13–15). Loss of KIF18A in multiple cell lines with CIN lead to decreased proliferation, multipolar spindles, and cell death (10–12). Importantly, these effects were not observed upon KIF18A knockdown in diploid cell lines (10–12). These results suggest that KIF18A is a promising therapeutic target for CIN cells.

Identifying molecular regions of KIF18A to target via small molecule inhibitors will be an important step in pursuing KIF18A inhibition as a therapeutic strategy. KIF18A activity is normally regulated during mitosis via binding to Kinesin Family Binding Protein (KIFBP) (16). KIFBP inhibits the activities of KIF18A and a subset of other kinesins by altering the conformation of the conserved alpha-4 helix within the motor domain, which in turn reduces the interaction of kinesins with the microtubule (17,18). A novel series of KIF18A inhibitors, which are currently being tested in clinical trials, have also been predicted to contact the alpha-4 helix region of KIF18A (19). Thus, the alpha-4 helix may represent an important site for regulating KIF18A activity. Consistent with this, a serine residue within alpha-4 helix (S284) has been identified as a phospho-residue via mass spectrometry (20). Here, we demonstrate that modulation of KIF18A activity through S284 mutations or chemical inhibition causes similar relocalization of the protein within the spindle and reduces the proliferation of CIN cells. These results confirm that multiple CIN tumor types depend on KIF18A activity and provide support for targeting KIF18A’s alpha-4 helix region as treatment strategy for chromosomally unstable tumors.

## RESULTS

### Alteration at Serine 284 in the highly conserved alpha-4 helix results in accumulation of KIF18A at spindle poles

Alpha-4 helix in the motor domain of KIF18A is highly conserved across various kinesin families and KIF18A homologs (Figure 1A). The helix is known to play a role in microtubule binding and nucleotide gating (21), facilitating the ability of the motor to move towards the plus ends of microtubules. Mass spectrometry data indicate that serine 284 (S284) within alpha-4 helix is phosphorylated in cells, suggesting that KIF18A activity may be regulated via alpha-4 helix phosphorylation (20). To investigate the effects of charge changes at S284, we generated HeLa Kyoto cell lines that inducibly express GFP-tagged wild-type KIF18A, phospho-null KIF18A (S284A), or a phosphomimetic KIF18A (S284D). Each construct contained siRNA-resistant, silent mutations and expression was induced following treatment with either control or KIF18A-specific siRNAs. Quantification of KIF18A immunofluorescence indicates that wild-type and mutant KIF18A constructs were expressed at similar levels (Figure S1). Analysis of KIF18A localization revealed that both S284A and S284D KIF18A mutants result in accumulation of KIF18A protein near spindle poles rather than at kinetochore microtubule plus ends (Figure 1B-C). This altered localization is unique compared to other reported KIF18A mutants that are predicted to compromise microtubule binding, suggesting that mutation of S284 may alter KIF18A activity through a mechanism that does not inhibit interaction with microtubules (14,22,23). The pole localization was even more apparent in live cells expressing GFP-KIF18A constructs, where the S284D mutant had a more pronounced effect, with less evidence of motor accumulation on the mitotic spindle compared to the S284A mutant (Figure 1D).

**Figure 1:**
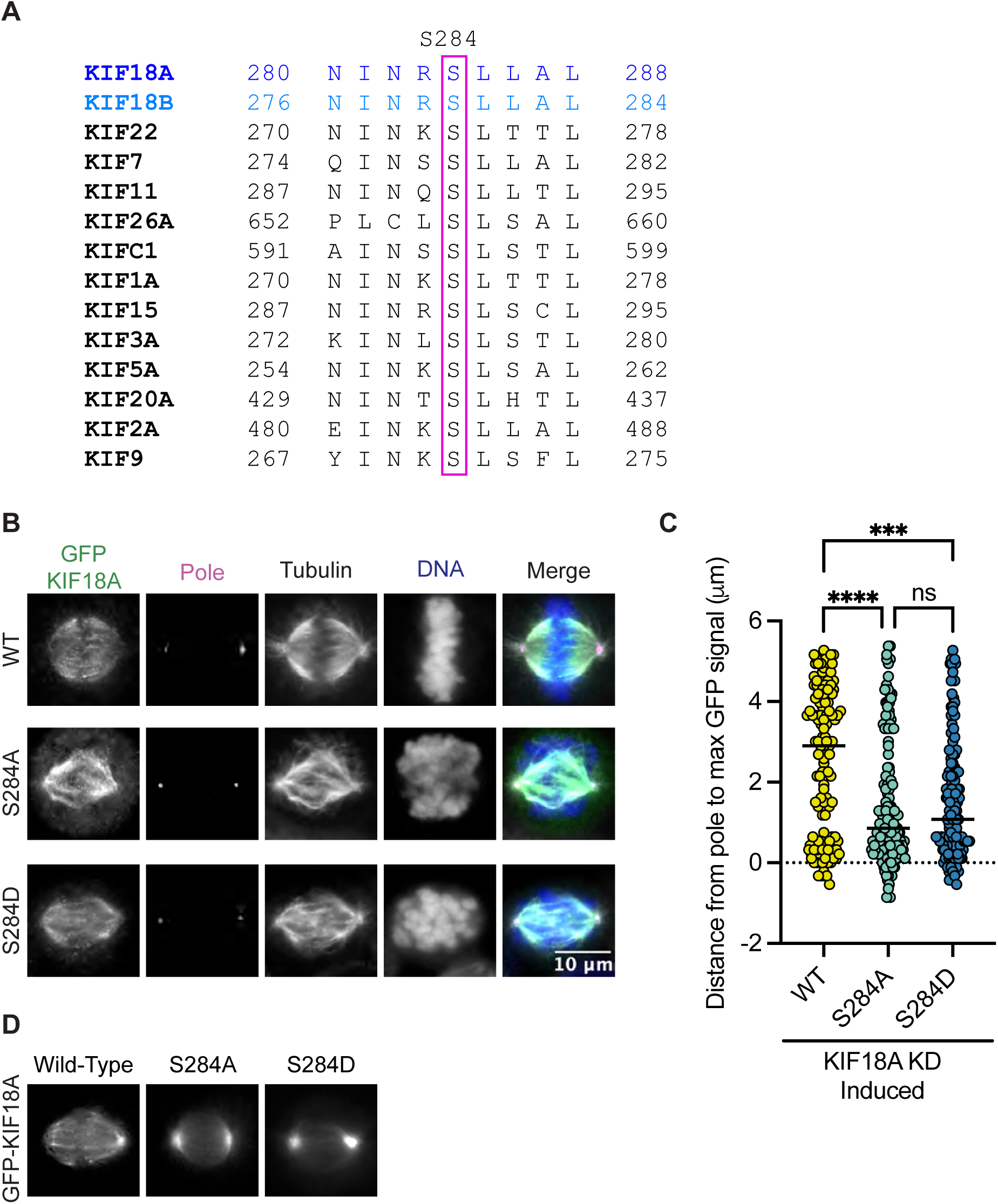
Mutation of KIF18A S284 alters its localization. **(A)** Sequence alignment of the alpha-4 region from kinesin family proteins. **(B)** Immunofluorescence images of HeLa Kyoto GFP-KIF18A inducible cells fixed and stained 24 hours after endogenous KIF18A knockdown and induction of indicated GFP-KIF18A construct. Scale bar = 2 μm. Text colors indicate pseudo color in merged image. From left to right, KIF18A antibody staining, γ-Tubulin antibody staining, α-Tubulin antibody staining, DNA/DAPI, merged image. **(C)** Quantification of distance from pole to maximum GFP-KIF18A signal in HeLa Kyoto cells using line scan analysis. Fluorescence values were normalized and aligned to the peak pole (γ-Tubulin) signal intensity. Solid line indicates median. Each dot represents an individual line scan. HeLa Kyoto GFP-KIF18A WT = 136 line scans, GFP-S284D = 130 line scans, and GFP-S284A = 144 line scans. Data acquired from three experimental replicates. Statistical results shown for a Kruskal-Wallis with Dunn’s Multiple Comparisons test; P values: < 0.05 (*), < 0.01 (**), < 0.001 (***), <0.0001 (****). N.s. indicates not significant (> 0.05).

### Mutation of S284 leads to loss of KIF18A function

KIF18A knockdown (KD) leads to unaligned chromosomes and increased spindle length (13,14). To investigate if mutations at S284 compromise KIF18A function during mitosis, we treated cells with siRNAs targeting KIF18A and induced expression of wild-type, S284A, or S284D KIF18A containing silent mutations that confer siRNA-resistance. KIF18A function was assessed by measuring chromosome alignment and spindle length in late prometaphase and metaphase cells. We determined the full width at half maximum of the chromosome distribution within an individual cell as a metric of chromosome alignment (24,25). Wild-type KIF18A restored chromosome distribution to levels seen in control KD cells, however, the S284A and S284D mutants were unable to promote chromosome alignment (Figure 2A-B). Additionally, while wild-type KIF18A expression resulted in spindle lengths comparable to those in control cells, neither S284A nor S284D mutants were able to properly regulate spindle length (Figure 2C). This work indicates that mutation of S284 in alpha-helix 4 results in loss of KIF18A function.

**Figure 2:**
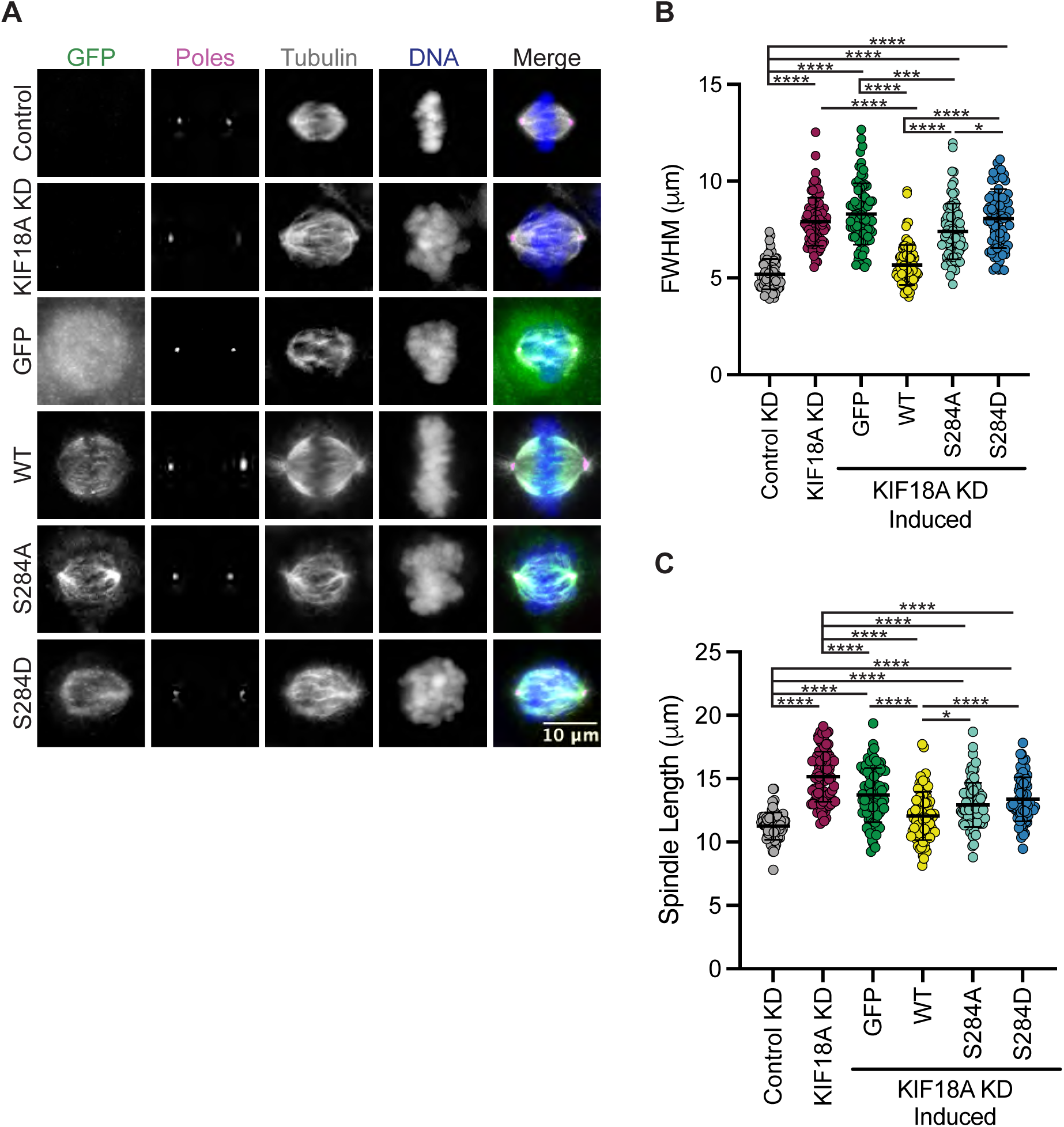
Expression of KIF18A S284 does not rescue KIF18A knockdown phenotypes. **(A)** Immunofluorescence images of HeLa Kyoto GFP KIF18A inducible cells fixed and stained 24 hours after endogenous KIF18A knockdown and induction of respective GFP KIF18A construct. Scale bar 10 μm. Colors indicate pseudo color in merged image. From left to right GFP antibody staining, poles/γ-Tubulin antibody staining, α-Tubulin antibody staining, DNA/DAPI, merged image. **(B)** Plot of FWHM (full-width half maximum) values measured for chromosome distributions in HeLa Kyoto GFP KIF18A inducible cells. Solid horizontal line indicates mean, vertical lines indicate standard deviation. Each dot represents a single cell. Data acquired from three experimental replicates. A one-way ANOVA with Tukey’s test for multiple comparisons was run, P-value: < 0.05 (*), < 0.01 (**), < 0.001 (***), <0.0001 (****). If no significance is indicated, differences wereas not significant (> 0.05). **(C)** Quantification of spindle length in HeLa Kyoto GFP KIF18A inducible cells. Solid horizontal line indicates mean vertical lines indicate standard deviation. Each dot represents a single cell. The following cell numbers were analyzed for each condition in B and C: control siRNA= 75, KIF18A siRNA = 84, GFP = 80, GFP-KIF18A WT = 81, GFP-S284A = 83, GFP-S284D = 78. Data acquired from three experimental replicates. A one-way ANOVA with Tukey’s test for multiple comparisons was run. P value style: < 0.05 (*), < 0.01 (**), < 0.001 (***), <0.0001 (****). If no significance is indicates result was not significant (> 0.05).

### KIF18A S284D causes mitotic arrest and multipolar mitotic spindles

Chromosomally unstable cell lines depend on KIF18A for mitotic progression and maintenance of bipolar mitotic spindles (10–12). To determine if KIF18A S284 mutants are able to promote normal mitotic progression, we determined the percentage of HeLa cells in mitosis following KIF18A KD and expression of S284 mutants. While KIF18A-S284A was able to promote mitotic progression to a greater extent than KIF18A-S284D, both mutants resulted in a significant increase in the percentage of mitotic cells compared to controls (Figure 3A-B). These data indicate that cells expressing KIF18A-S284D experience delays in mitotic progression similar to those that occur following KIF18A depletion (13,14,26).

**Figure 3:**
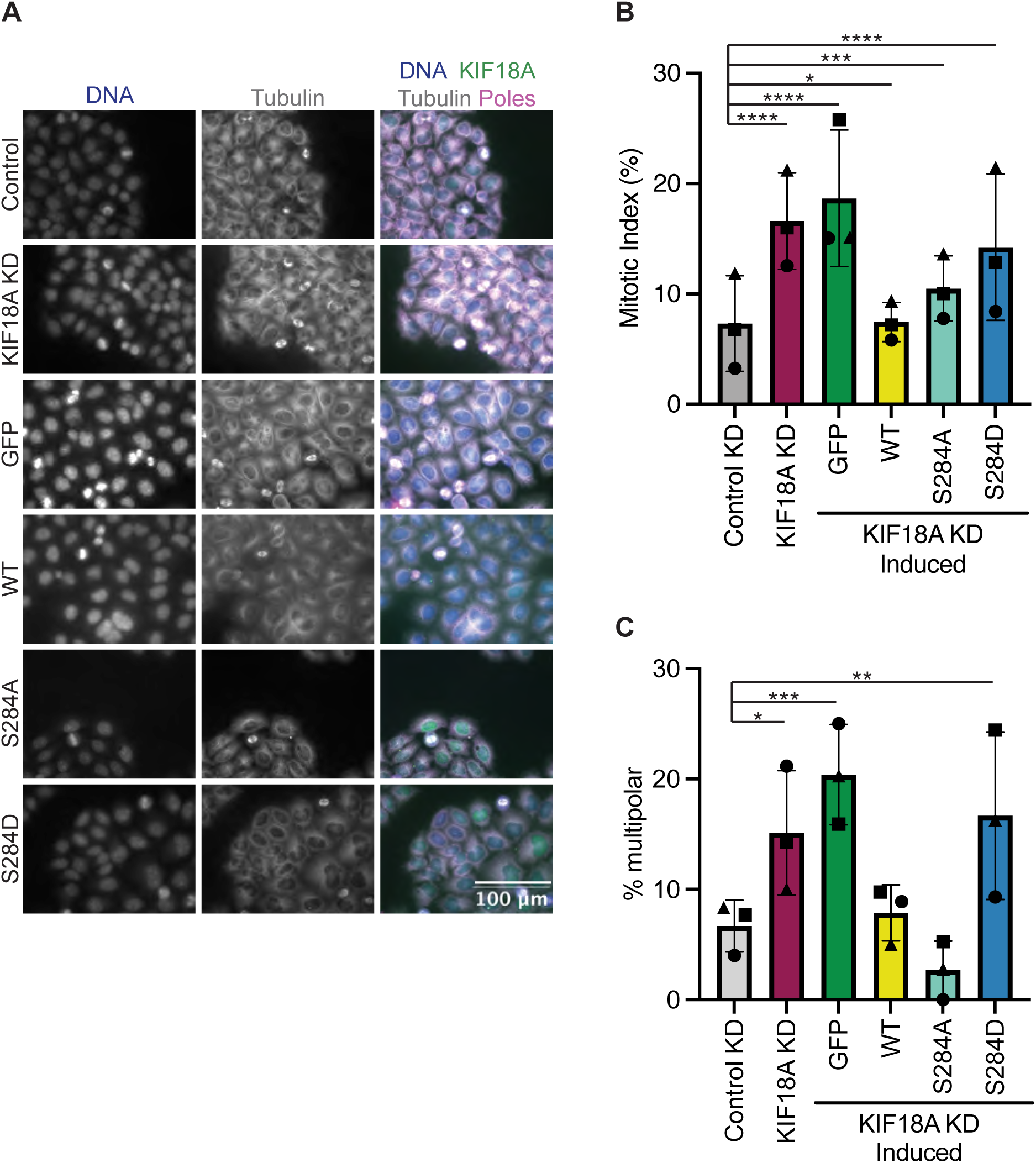
Mutation of KIF18A S284 to S284D leads to mitotic arrest and multipolar spindles. **(A)** Immunofluorescence images of HeLa Kyoto GFP KIF18A inducible cells fixed and stained 24 hours after endogenous KIF18A knockdown and induction of respective GFP KIF18A construct. Scale bar 100 μm. Colors indicate pseudo color in merged image. From left to right DNA/DAPI, α-Tubulin antibody staining, merged image. Merged image includes KIF18A and spindle pole antibody staining in addition to DNA and α-Tubulin antibody staining. **(B)** Quantification of mitotic index (% cells in mitosis) in HeLa Kyoto GFP KIF18A inducible cell lines. Bars indicate overall mean, circle indicates mean from experimental replicate one, square indicates mean from experimental replicate two, and the triangle indicates the mean from experimental replicate three. Total number of cells analyzed for HeLa Kyoto KIF18A KD = 1402 cells, HeLa Kyoto GFP = 997 cells, HeLa Kyoto wild type = 2136 cells, HeLa Kyoto S284A = 1364 cells, and HeLa Kyoto S284D = 1574 cells. **(C)** Quantification of multipolarity (% multipolar cells) in HeLa Kyoto GFP KIF18A inducible cell lines. Bars indicate overall mean, circle indicates mean from experimental replicate one, square indicates mean from experimental replicate two, and the triangle indicates the mean from experimental replicate three. Total number of cells analyzed for HeLa Kyoto: KIF18A KD = 1402, GFP = 997, GFP-KIF18A WT = 2136, GFP-S284A = 1364, and GFP-S284D = 1574.

Loss of KIF18A function in chromosomally unstable cells also results in an increase in the number of mitotic cells with multipolar spindles (11). To determine if S284 mutants can promote spindle bipolarity, the percentage of multipolar mitotic cells was determined following KIF18A KD and expression of wild-type or S284 mutant KIF18A. While wild-type and S284A KIF18A expressing cells formed multipolar spindles at a level similar to controls, KIF18A-S284D expressing cells displayed a significant increase in multipolar spindles, similar to levels seen following KIF18A KD alone (Figure 3C).

### KIF18A-S284D expressing cells display reduced proliferation

Mitotic arrest and multipolar spindles in KIF18A-depleted, chromosomally unstable tumor cells strongly correlate with decreased proliferation (11). To probe if expression of KIF18A-S284 mutants also result in proliferation defects, we performed a microscopy-based kinetic proliferation assay in cells treated with KIF18A siRNAs expressing wild type or S284 mutant KIF18A (11). Cell populations expressing GFP alone or KIF18A-S284D displayed significantly lower proliferation during a 5-day time-course than cells expressing wild type or KIF18A-S284A (Figure 4A-C). The observed decrease in proliferation for KIF18A-S284D correlates with the measured effects of this mutant on mitotic arrest and spindle multipolarity.

**Figure 4:**
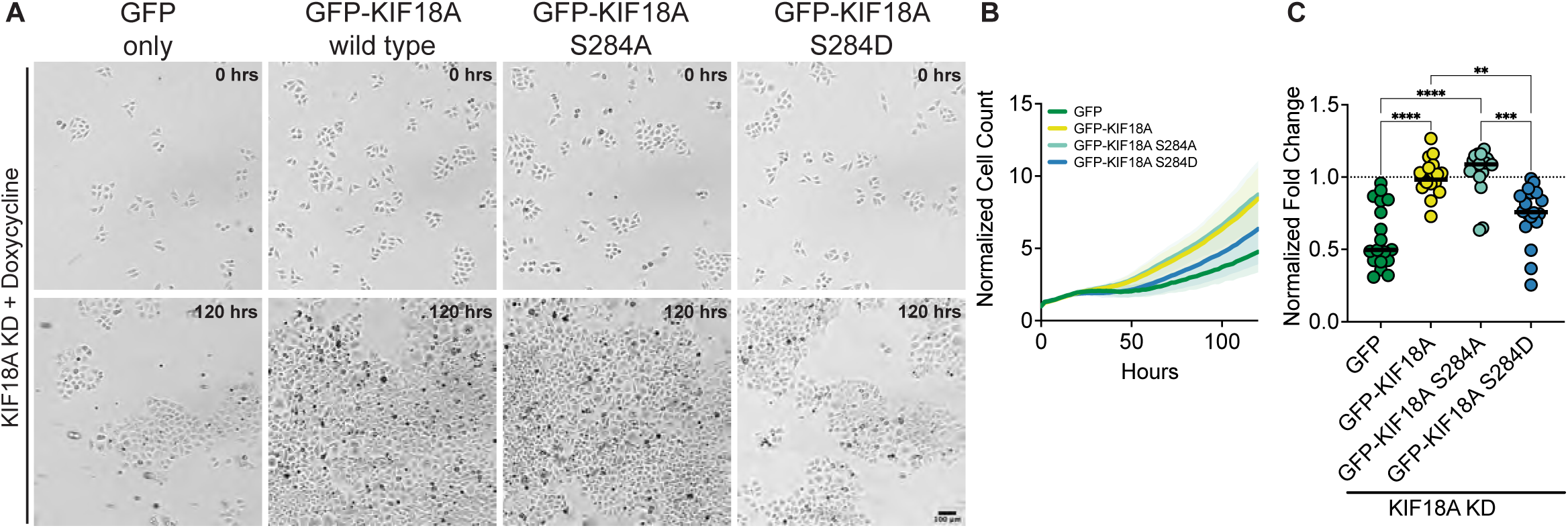
Phospho-mimetic mutation of KIF18A S284 leads to decreased proliferation of HeLa Kyoto cells. **(A)** Brightfield images of HeLa Kyoto cells from time-lapse proliferation assays after knockdown of endogenous KIF18A and induced expression of indicated construct. Scale bar 100 μm. (Top) 0 hrs. (Bottom) 120 hours. **(B)** Normalized Cell Count versus Time (hours) from automated bright field imaging. The cell counts from each time point were normalized to the number of cells at the initial time point. Lines represent means and shaded areas are standard deviation. Data shown from three experimental replicates. **(C)** Normalized Fold Change after 120 hours for each condition. All conditions were normalized to GFP-KIF18A wild type rescue, indicated by dotted line. Each dot represents an individual well of cells. Data shown from three experimental replicates. Statistical results are shown for a Kruskal-Wallis with Dunn’s Multiple Comparisons test. P value style: < 0.05 (*), < 0.01 (**), < 0.001 (***), <0.0001 (****). N.s. indicates not significant (> 0.05).

### Chemical inhibition of KIF18A with Compound 3 or Sovilnesib phenocopies KIF18A knockdown phenotypes

Interestingly, S284 in KIF18A is located adjacent to residues within the alpha-4 helix that are proposed to form a binding pocket for a series of recently described KIF18A inhibitors (19). One of these inhibitors, Sovilnesib, is currently being tested in clinical trials (Identifier NCT04293094) for patients with advanced p53-mutated tumors, highlighting the therapeutic potential of KIF18A-targeting drugs. If these inhibitors inactivate KIF18A through effects on alpha-4, we might expect the specific phenotypes caused by the expression of KIF18A-S284 mutants and inhibitor treatment to be similar. To address this question, we synthesized Sovilnesib, as well as a similar derivative with a single heteroatom substitution (19), hereafter referred to as Compound 3 (Figure 5A). Biochemical studies *in vitro* with purified KIF18A revealed a half-maximal inhibitory concentration (IC_50_) of 8.2 nanomolar (nM) for Compound 3 and 41.3 nM for Sovilnesib (Figure 5B). These results confirm that Compound 3 and Solvilnesib are potent inhibitors of KIF18A’s microtubule-stimulated ATPase activity.

**Figure 5:**
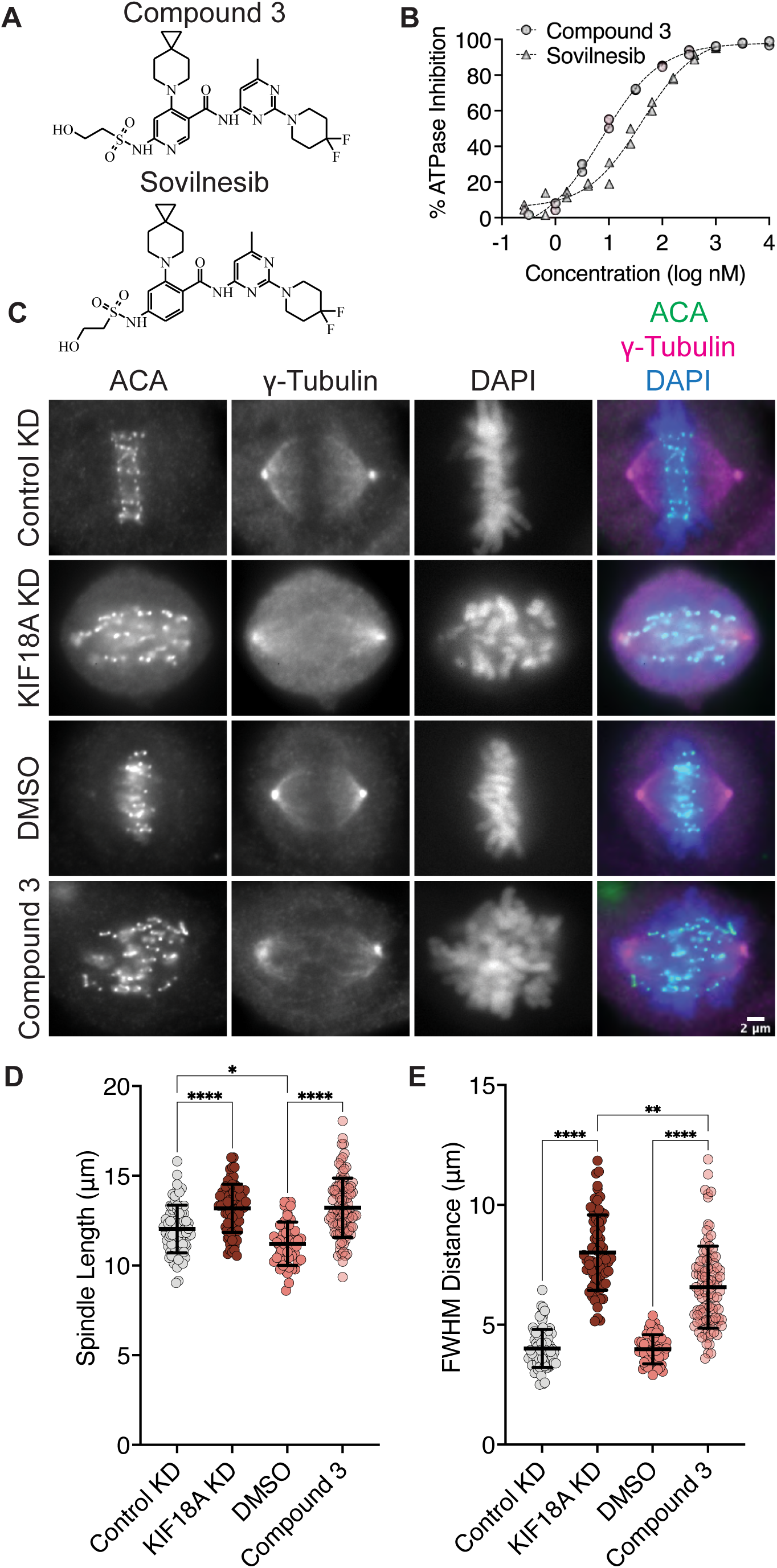
KIF18A inhibitors mimic KIF18A knockdown phenotypes in MDA-MB-231 cells. **(A)** Chemical structure of Compound 3. **(B)** % Inhibition of KIF18A microtubule stimulated ATPase activity *in vitro* versus logarithmic concentration of the indicated KIF18A inhibitors. Representative data from two experimental replicates of each is shown. Dashed lines indicate non-linear fit of the data. **(C)** Immunofluorescence images of MDA-MB-231 cells fixed and stained after 24 hours of indicated treatments. Scale bar 2 μm. **(D)** Graph of spindle lengths measured in cells fixed and stained 24 hours after the indicated treatments. Each dot represents a single cell. Mean ± standard deviation is displayed. Statistical results are shown for a Kruskal-Wallis with Dunn’s Multiple Comparisons test. P value style: < 0.05 (*), < 0.01 (**), < 0.001 (***), <0.0001 (****). N.s. indicates not significant (> 0.05). **(E)** Graph of full-width at half maximum (FWHM) of centromere fluorescence distribution along the length of the spindle measured in cells fixed and stained 24 hours after the indicated treatments. Each dot represents a single cell. Mean ± standard deviation is displayed. Statistical results are shown for a Kruskal-Wallis with Dunn’s Multiple Comparisons test. P value style: < 0.05 (*), < 0.01 (**), < 0.001 (***), <0.0001 (****). n.s. indicates not significant (> 0.05). The following cell numbers were analyzed for **(D-E)**: Control siRNA = 76, KIF18A siRNA = 69, DMSO = 51, Compound 3 = 96.

To determine whether these compounds inhibit KIF18A in cells, we treated chromosomally unstable triple negative breast cancer (TNBC) MDA-MB-231 cells for 24 hours with dimethyl sulfoxide (DMSO), 250 nM Compound 3 or Sovilnesib, Control siRNA, or KIF18A siRNA and measured chromosome alignment and spindle length (Figure 5C-E and Figure S2). Treatment of cells with KIF18A siRNA or 250 nM KIF18A inhibitor resulted in increased spindle length (Figure 5D and Figure S2A) as compared to Control siRNA and DMSO treatment, respectively. Similarly, treatment of MDA-MB-231 cells with 250 nM KIF18A inhibitor lead to defects in chromosome alignment, though to a lesser extent than KIF18A KD (Figure 5E and Figure S2B). The difference in the magnitude of the shift in chromosome alignment between Compound 3 and KIF18A siRNA treatment may represent biological differences in inhibition versus KD due to the continued presence of KIF18A protein. Together, these results suggest that chemical inhibition of KIF18A by Compound 3 mimics KIF18A siRNA KD phenotypes in MDA-MB-231 cells.

### KIF18A accumulates at spindle poles following inhibitor treatment

One defining feature of the KIF18A-S284 mutants is their accumulation at spindle poles when expressed in mitotic cells (Figure 1B). Normally, KIF18A accumulates at the plus-ends of microtubules, where it regulates chromosome movements and promotes chromosome alignment during metaphase (13,14,25). To determine if KIF18A inhibitor treatment leads to similar aberrant KIF18A localization away from microtubule plus-ends, we measured KIF18A localization in multiple cell types after treatment with 250 nM Compound 3, Sovilnesib, or DMSO control. To confirm changes in KIF18A localization in both chromosomally stable and unstable cell lines, we used diploid retinal pigment epithelial-1 (RPE1) cells and two chromosomally unstable cell lines, colorectal adenocarcinoma HT-29 cells and MDA-MB-231 cells. In all cell types, we observed a shift in localization of KIF18A away from microtubule plus-ends toward the spindle poles (Figure 6A-F and Figure S3). To quantify this change in localization, line scan analyses were conducted to determine the distance from the spindle pole to the max KIF18A signal in each half spindle. Cells treated with DMSO had characteristic accumulation of KIF18A toward the plus-ends of microtubules several microns away from the pole, whereas cells treated with 250 nM Compound 3 or Sovilnesib exhibited accumulation of KIF18A in the vicinity of the spindle poles (Figure 6A-F and Figure S3 A-B). These results demonstrate that KIF18A inhibitor treatment leads to dynamic accumulation of KIF18A at spindle poles in both chromosomally stable and unstable cell types.

**Figure 6:**
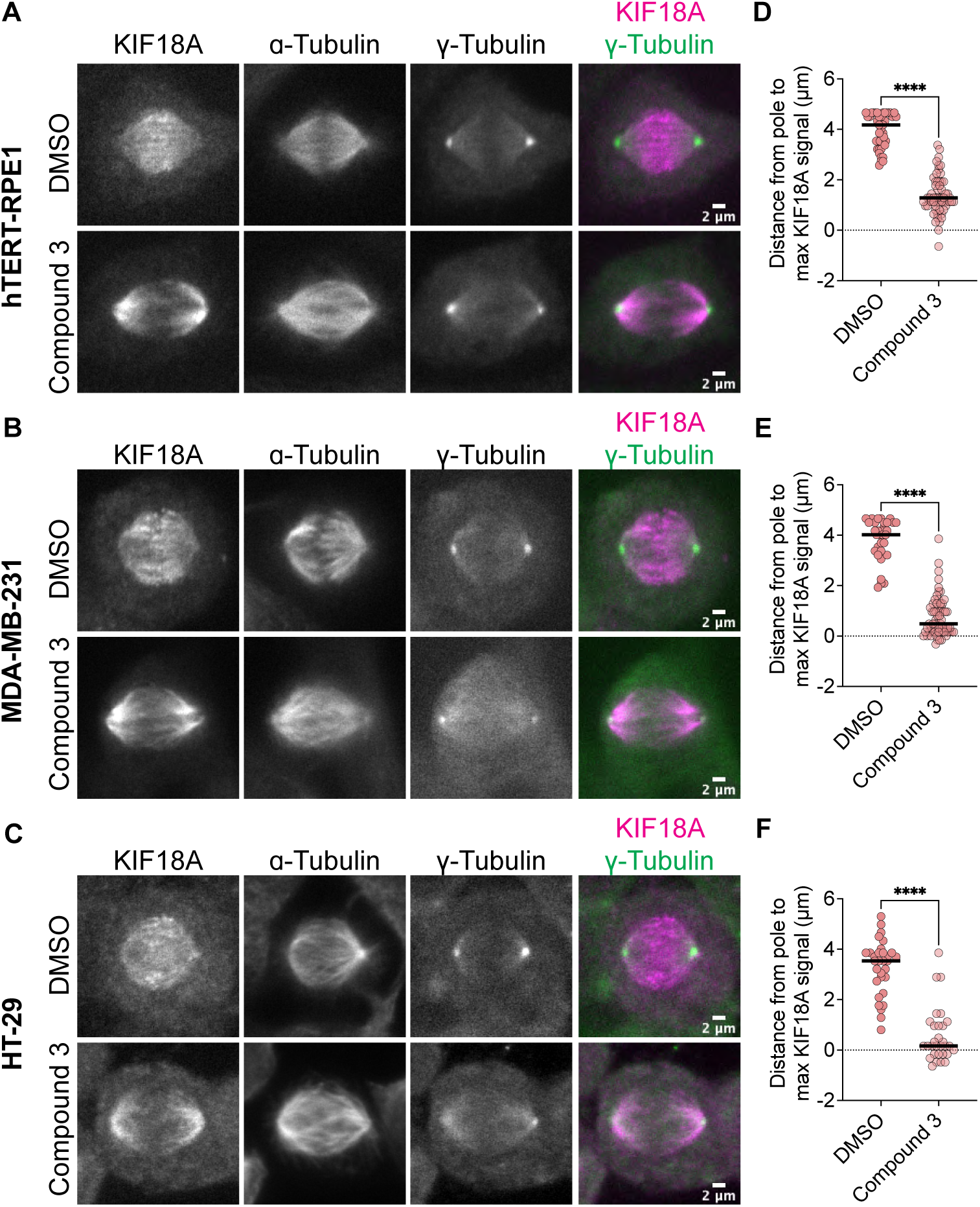
Compound 3 disrupts microtubule plus-end localization of KIF18A. **(A-C)** Immunofluorescence images of hTERT-RPE1 **(A)**, MDA-MB-231 **(B)**, and HT-29 **(C)** cells fixed and stained 24 hours after DMSO or 250 nM Compound 3 treatment to visualize KIF18A localization. Left to right: KIF18A antibody staining, α-Tubulin antibody staining, γ-Tubulin antibody staining, merge of KIF18A (magenta) and γ-Tubulin (green) staining. Scale bar 2 μm. **(D-F)** Plots of distance from max KIF18A signal to the spindle pole derived from line scan analyses of KIF18A distribution in in the indicated cell types. Each dot represents a single cell. Dark bars represent the mean values. Statistical results are shown for a Kruskal-Wallis with Dunn’s Multiple Comparisons test. P value: < 0.05 (*), < 0.01 (**), < 0.001 (***), <0.0001 (****). The following number of line scans were analyzed for the indicated cell lines and conditions: HT-29 DMSO = 31 line scans, HT-29 Compound 3 = 30 line scans, MDA-MB-231 DMSO = 33 line scans, MDA-MB-231 Compound 3 = 59 line scans, RPE1 DMSO = 42 line scans, RPE1 Compound 3 = 59 line scans.

### KIF18A inhibitors cause mitotic arrest and reduced proliferation of chromosomally unstable cells

Given that KIF18A inhibitor treatment mimics the chromosome alignment and spindle length phenotypes observed following KIF18A KD, we predicted that inhibition of KIF18A with Compound 3 or Sovilnesib would lead to mitotic arrest in chromosomally unstable cell lines but not diploid cells, as previously observed (10–12,26). To test this prediction, we measured mitotic index in diploid RPE1 cells and the chromosomally unstable cell lines MDA-MB-231 and HT-29. In diploid RPE1 cells, KIF18A inhibitor treatment and KIF18A KD did not affect the mitotic index as compared to DMSO or Control KD (Figure 8A-B and Figure S4A). In contrast, MDA-MB-231 and HT-29 cells treated with KIF18A inhibitors or KIF18A siRNAs displayed a significant increase in the mitotic index (Figure 8C-F and Figure S4B-C). Furthermore, MDA-MB-231 and HT-29 cells displayed a significant increase in multipolar spindles following KIF18A inhibitor treatment, but RPE1 cells did not (Figure 8G-H and Figure S4D-F). Of note, the HT-29 cells treated with KIF18A inhibitors resulted in a higher percentage of multipolar spindles than bipolar spindles (Figure 8H and Figure S4F). These results suggest that both chemical inhibition of KIF18A and KIF18A KD result in mitotic arrest and abnormal spindle formation in chromosomally unstable cell lines, but not diploid cells.

**Figure 7:**
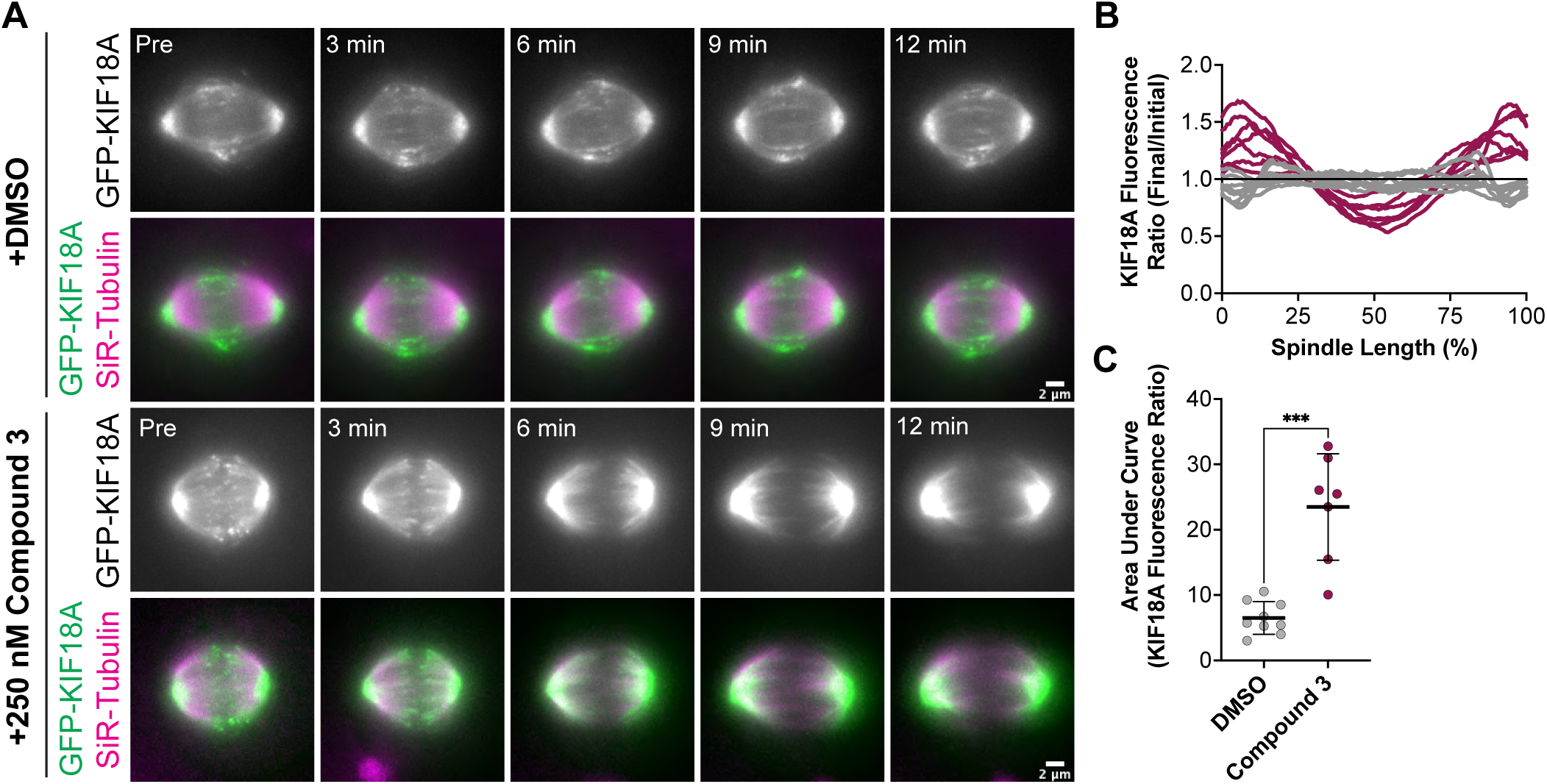
Compound 3 leads to rapid relocalization of KIF18A to the poles. **(A-B)** Time-lapse images of hTERT-RPE1 cells inducibly expressing wild type GFP-KIF18A after knockdown of endogenous KIF18A. Left: GFP-KIF18A. Middle: SiR-Tubulin. Right: Merge of GFP-KIF18A (green) and SiR-Tubulin (magenta). **(A)** Indicated timepoints from time-lapse images of hTERT-RPE1 cells before and after DMSO addition. Scale bar 2 μm. **(B)** Indicated timepoints from time-lapse images of hTERT-RPE1 cells before and after 250 nM Compound 3 addition. Scale bar 2 μm. **(C)** Graph of GFP-KIF18A fluorescence signal (Final Fluorescence Values divided by Initial Fluorescence Values from region of interest capturing the entire spindle) versus % Spindle Length (Length of individual spindles were normalized on a scale from 0 to 100% for cell-to-cell comparison). Gray lines are DMSO treated cells, magenta lines are 250 nM Compound 3 treated cells. Each line represents an individual cell. Data compiled from three experimental replicates. **(D)** Quantification of area under the curve from graph in **(C)**. Bars are mean +/- standard deviation. Dots represent individual cells from 3 independent experiments (N=9 for DMSO, and N=7 for Compound 3). Statistical results displayed from a Mann-Whitney test. P value style: < 0.05 (*), < 0.01 (**), < 0.001 (***), <0.0001 (****). n.s. indicates not significant (> 0.05).

**Figure 8:**
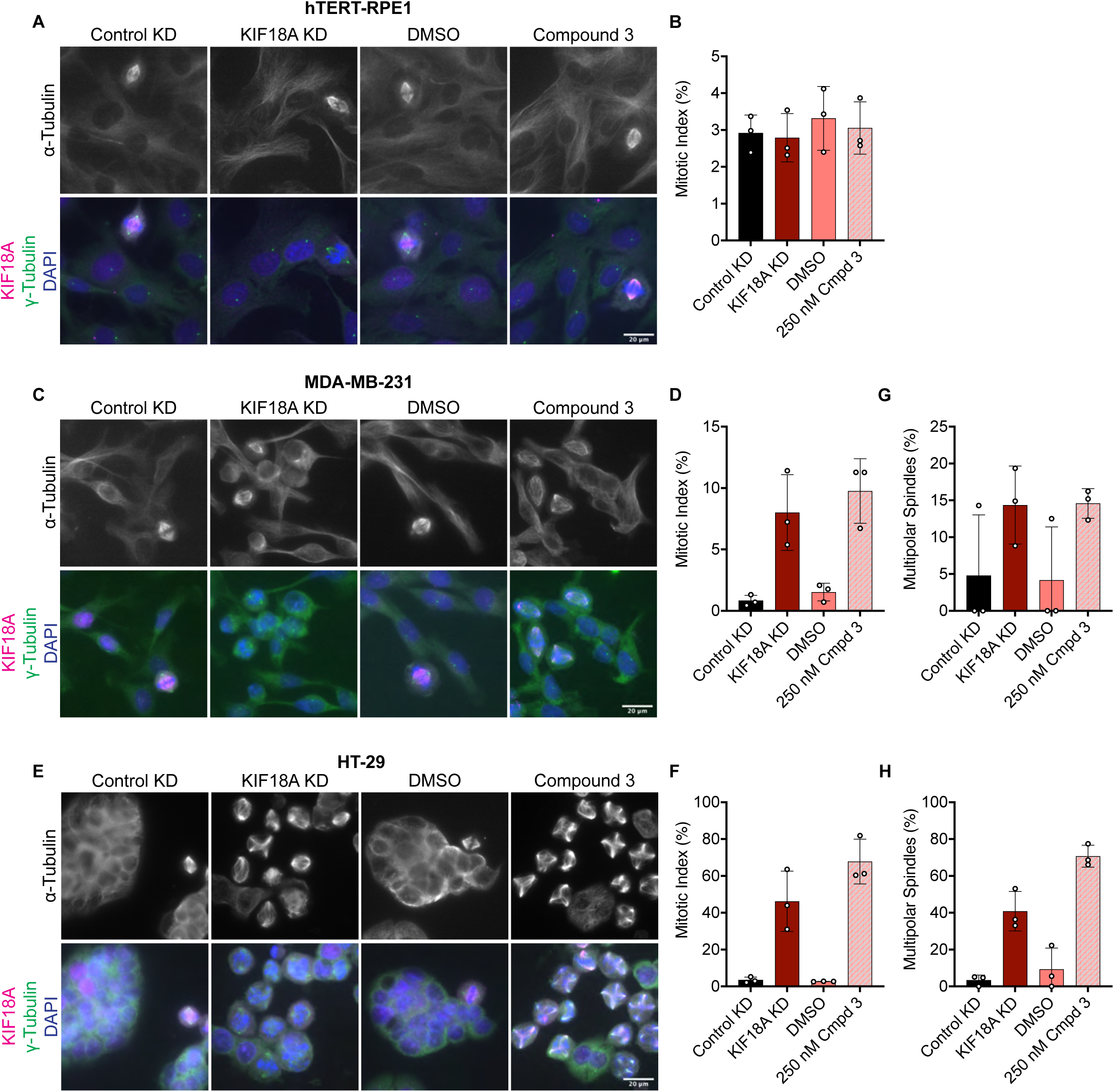
Compound 3 treatment leads to mitotic arrest and multipolar spindles in chromosomally unstable breast and colorectal cancer cells. **(A)** Immunofluorescence images of hTERT-RPE1 cells fixed and stained 24 hours after indicated treatments. Scale bar 20 μm. Top: α-Tubulin only. Bottom: Merge of γ-Tubulin (green), KIF18A (magenta), and DAPI/DNA (blue). **(B)** Quantification of mitotic index (% of total cells in mitosis) in hTERT-RPE1 cells 24 hours after indicated treatments. Bars are mean +/- standard deviation. Each dot indicates an experimental replicate. **(C)** Immunofluorescence images of MDA-MB-231 cells fixed and stained 24 hours after indicated treatments. Scale bar 20 μm. Top: α-Tubulin only. Bottom: Merge of γ-Tubulin (green), KIF18A (magenta), and DAPI/DNA (blue). **(D)** Quantification of mitotic index (% of total cells in mitosis) in MDA-MB-231 cells 24 hours after indicated treatments. Bars are mean +/- standard deviation. Each dot indicates an experimental replicate. **(E)** Immunofluorescence images of HT-29 cells fixed and stained 24 hours after indicated treatments. Scale bar 20 μm. Top: α-Tubulin only. Bottom: Merge of γ-Tubulin (green), KIF18A (magenta), and DAPI/DNA (blue). **(F)** Quantification of mitotic index (% of total cells in mitosis) in HT-29 cells 24 hours after indicated treatments. Bars are mean ± standard deviation. Each dot indicates an experimental replicate. **(G-H)** Quantification of multipolar spindles (% of total spindles) in MDA-MB-231 cells **(G)** and HT-29 cells **(H)** 24 hours after indicated treatments. Bars are mean +/- standard deviation. Each dot indicates an experimental replicate. The following total cell numbers were analyzed for each condition and cell line: HT-29: Control siRNA = 2257, KIF18A siRNA = 1319, DMSO = 2014, Compound 3 = 935 cells; MDA-MB-231: Control siRNA = 2298, KIF18A siRNA = 1928, DMSO = 2468, Compound 3 = 2115; RPE1: Control siRNA = 871, KIF18A siRNA = 940, DMSO = 1052, Compound 3 = 1049; HeLa Kyoto Control siRNA = 1558, siRNA = 1402, GFP = 997, GFP-KIF18A WT = 2136, GFP-S284A = 1364, and GFP-S284D = 1574.

To determine whether the mitotic arrest phenotypes observed after KIF18A inhibitor treatment also lead to a decrease in growth, we measured the proliferation of diploid RPE1 and chromosomally unstable MDA-MB-231 and HT-29 cells in increasing concentrations of Compound 3 or Sovilnesib using a kinetic proliferation assay (11). The proliferation of diploid RPE1 cells was unaffected by the KIF18A inhibitors up to 1 μM (Figure 9A-C and Figure S5A-B). In contrast, the proliferation of MDA-MB-231 and HT-29 cells was significantly reduced with increasing concentrations of KIF18A inhibitors (Figure 9D-I and Figure S5C-F). The increased sensitivity of the HT-29 cells to KIF18A inhibitors compared to MDA-MB-231 cells is consistent with our previous findings that the fold decrease in proliferation was inversely correlated with the fold increase in multipolar spindles following KIF18A KD (11). Furthermore, chemical inhibition of KIF18A specifically decreased proliferation of chromosomally unstable cell lines but did not affect the growth of diploid cells, highlighting the therapeutic potential of KIF18A inhibitors for specifically targeting tumor cells.

**Figure 9:**
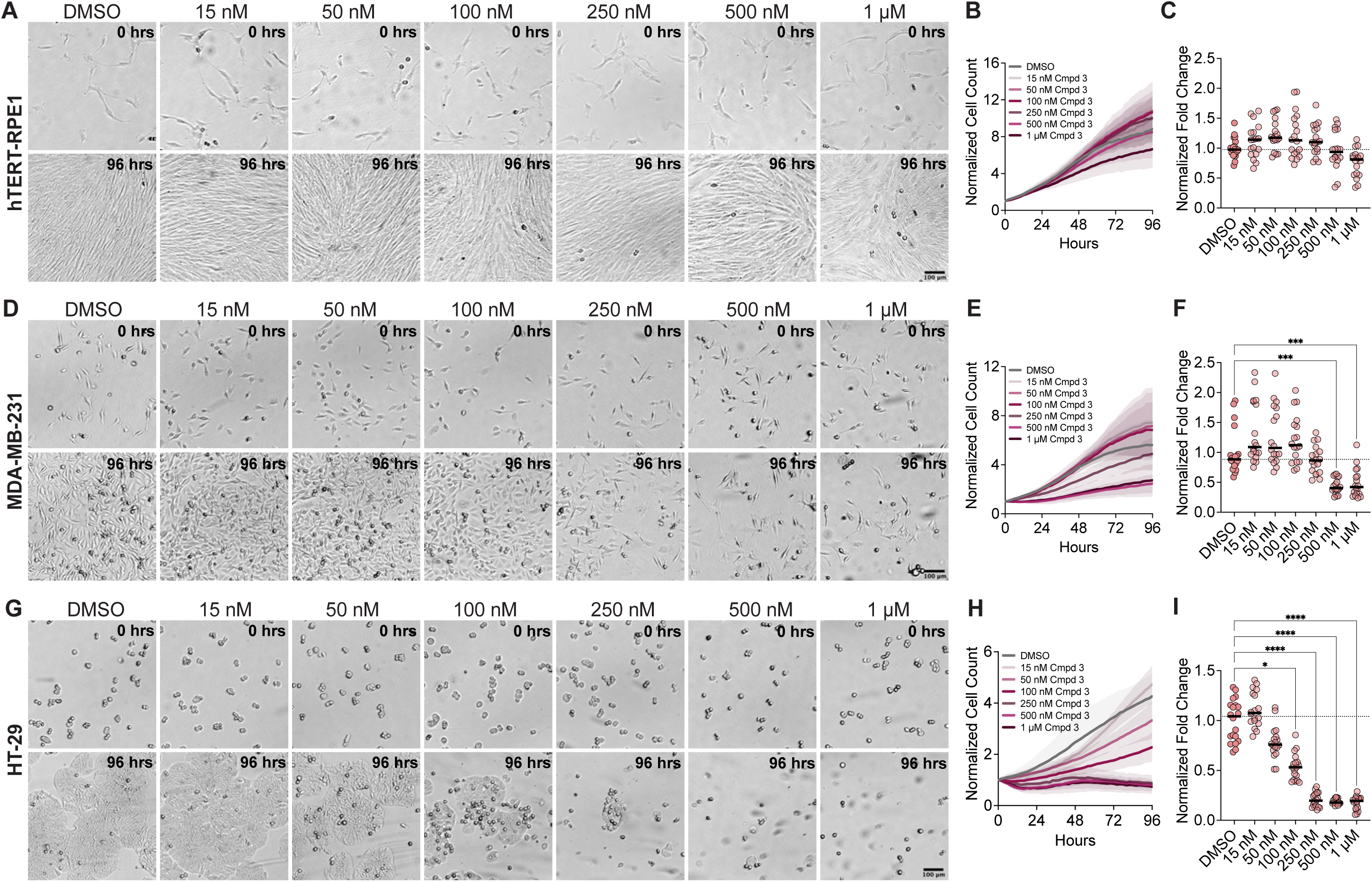
Compound 3 reduces the proliferation of chromosomally unstable cells in a dose-dependent manner. **(A)** Bright field images of hTERT-RPE1 cells from time-lapse proliferation assays with the indicated concentrations of Compound 3. Scale bar 100 μm. (Top) 0 hrs. (Bottom) 96 hours. **(B)** Plot of normalized cell count over time for cells treated with the indicated concentrations of Compound 3. Lines indicate mean and shaded area denotes SD. **(C)** Plot of normalized hTERT-RPE1 cell count (displayed as a % of DMSO control) as a function of Compound 3 concentration. **(D)** Bright Field images of MDA-MB-231 cells from time-lapse proliferation assays with the indicated concentrations of Compound 3. Scale bar 100 μm. (Top) 0 hrs. (Bottom) 96 hours. **(E)** Plot of normalized cell count over time for MDA-MB-231 cells treated with the indicated concentrations of Compound 3. Lines indicate mean and shaded area denotes SD. **(F)** Plot of normalized MDA-MB-231 cell count (displayed as a % of DMSO control) as a function of Compound 3 concentration. **(G)** Bright Field images of HT-29 cells from time-lapse proliferation assays with the indicated concentrations of Compound 3. Scale bar 100 μm. (Top) 0 hrs. (Bottom) 96 hours. **(H)** Plot of normalized cell count over time for HT-29 cells treated with the indicated concentrations of Compound 3. Lines indicate mean and shaded area denotes SD. **(I)** Plot of normalized HT-29 cell count (displayed as a % of DMSO control) as a function of Compound 3 concentration. For plots in **(C)**, **(F)**, and **(I)**, each dot indicates an individual well and bars indicate mean values. Statistical results displayed from a Mann-Whitney test. P value style: < 0.05 (*), < 0.01 (**), < 0.001 (***), <0.0001 (****). n.s. indicates not significant (> 0.05). Data shown on all plots are from three independent experiments.

## DISCUSSION

Here, we show that chemical inhibition of KIF18A and mutations that mimic phosphorylation of the alpha-4 helix lead to alterations in the localization of the motor away from the microtubule plus-ends toward the spindle poles. In addition to resulting in altered localization, both S284 mutations and chemical inhibition resulted in increased chromosome distribution and spindle length. Furthermore, chemical inhibition and S284D mutations resulted in defects in proliferation, an increased mitotic index, and increased multipolarity specifically in chromosomally unstable cells. These data further support KIF18A as a promising target for the treatment of CIN tumors.

The similar KIF18A localization pattern caused by chemical inhibition of KIF18A and S284 mutations suggests a possible mechanism of action whereby the chemical inhibitors induce alterations to the alpha-4 helix of KIF18A. This is consistent with modeling data suggesting that the KIF18A inhibitors tested in this study interact with a hydrophobic pocket of the KIF18A enzymatic domain between the alpha-4 and alpha-6 helices (19). Localization of KIF18A to spindle poles is different than the effects caused by mutations that are predicted to reduce KIF18A interaction with microtubules. In those cases, KIF18A is evenly distributed along microtubules and displays more cytoplasmic localization (14,22,23). Thus, we hypothesize that S284 mutations and the KIF18A inhibitors tested here may stabilize KIF18A in a conformation that is tightly bound to microtubules. This model is consistent with data indicating that compounds in this chemical series are more effective inhibitors of KIF18A ATPase activity in the presence of microtubules (19). In this case, the localization of KIF18A protein to spindle poles could be explained by microtubule flux within the spindle, where microtubule-associated KIF18A travels towards the minus-ends of microtubules as they are depolymerized (27,28). Consistent with this idea, a gap in KIF18A-GFP protein can be seen to expand from the center of the spindle towards the poles from 3-12 minutes after inhibitor treatment (Figure 7A). Thus, trapping KIF18A in a microtubule bound state may be an effective inhibition strategy for specifically reducing the proliferation of CIN tumor cells.

KIF18A is also required to maintain spindle bipolarity in CIN cells, which subsequently influences their dependence on KIF18A for proliferation (11). In this study, we show that both chemical inhibition of KIF18A and S284D mutations result in aberrant KIF18A localization to spindle poles and subsequent loss of spindle bipolarity. Interestingly, the S284A mutation does not have as pronounced of a localization defect and does not increase the percent of multipolar mitotic spindles. This lack of increase in multipolarity is correlated with a proliferation rate that more closely mimics that of cells expressing wild-type KIF18A, suggesting that the S284A mutant retains enough function to support mitotic progression. These data suggest KIF18A’s roles in maintaining spindle polarity and promoting progression through mitosis as being critical for CIN cell proliferation, consistent with previous observations (10–12). On the other hand, reduced chromosome alignment and longer spindle lengths in the S284A mutant expressing cells appear to be tolerable defects that do not result in an immediate reduction in proliferation. Similarly, diploid somatic cells can continue proliferating in the absence of KIF18A activity, despite the presence of chromosome alignment defects (22,26,29). In the case of the S284A mutant, we predict that enough KIF18A activity remains to support mitotic progression but not chromosome alignment. This potential graded phenotypic response following partial and more complete inhibition of KIF18A is useful to consider when evaluating whether KIF18A inhibitors produce effects that may potentially inhibit CIN tumor proliferation.

This work demonstrates that chemical inhibition of KIF18A closely mimics the differential effects of KIF18A KD in CIN and diploid cells in terms of single cell phenotypes and suggests the alpha-4 helix as a potential target for effective inhibition. Thus, our results support the idea that KIF18A inhibition may serve as a promising therapeutic treatment for CIN tumor types including breast, colorectal, cervical, and ovarian cancer. Future work will focus on determining how the alpha-4 helix is altered by mutations in S284 or binding to inhibitory compounds.

## MATERIALS AND METHODS

### Cell Culture and Transfections

HT-29 and MDA-MB-231 cells were purchased from ATCC. MDA-MB-231 and HT-29 cells were grown in DMEM/F12 medium (Gibco, 11320082) containing 10% FBS (Gibco, 16000044). HeLa Kyoto and RPE1 acceptor cell lines were both generous gifts from Ryoma Ohi, University of Michigan. HeLa Kyoto and RPE1 cells were cultured at 37°C with 5% CO_2_ in MEM-α medium (Gibco, 12561072) containing 10% Fetal Bovine Serum (FBS) (Gibco, 16000044). Acceptor cell lines were maintained in Blasticidin (Thermo Fisher Scientific, R21001) until generation of inducible clones, after which cell lines were maintained in Puromycin (Thermo Fisher Scientific, A11138-03). For siRNA transfections in a 24-well plate format, MDA-MB-231, HT-29, HeLa Kyoto and RPE1 cells were treated with 5 pmol siRNA that was preincubated for 5-10 minutes in Opti-MEM Reduced Serum Media (Gibco, 31985062) with Lipofectamine RNAiMax Transfection Reagent (Invitrogen, 13778150). KIF18A siRNAs used were a 1:1 mixture of two the following Silencer or Silencer Select Validated siRNAs (5’ to 3’ sequence): GCUGGAUUUCAUAAAGUGGtt (Ambion, AM51334), GCUUGUUCCAGAAUCGAGAtt (Ambion, 4390824) or CGUUAACUGCAGACGUAAAtt (Ambion, 4392420). Control siRNAs used were Ambion Silencer Select Negative Control #2 (Ambion, 4390846), Silencer Negative Control siRNA #2 (Ambion, AM4613), or Silencer Select Negative Control #1 (4390843).

### Generation and Validation of HeLa Kyoto and RPE1 Inducible Cell Lines

All KIF18A inducible cell lines were generated using previously described methods (26,30,31). Briefly a wild-type KIF18A siRNA and puromycin resistant plasmid was developed containing LoxP sites for recombination mediated cassette exchange. The LoxP containing plasmid was then transfected into HeLa Kyoto acceptor cells (32) and cells which had undergone recombination were selected for with puromycin. KIF18A wild-type(22), S284A (generated using mutagenesis), and S284D (generated through mutagenesis) siRNA resistant fragments, and pEM791 vector (32) were amplified with primers designed for Gibson Assembly (New England BioLabs). After confirming the correct sequence of the Gibson assembled plasmid, recombination was achieved by transfecting the acceptor cells with the KIF18A plasmid and recombinase using an LTX transfection (Thermo Fisher Scientific). Recombination was initially selected for with 1 μg/mL puromycin for 48 hours followed by a stricter selection with 2 μg/mL puromycin for 48 hours prior to switching back to 1 μg/mL puromycin for HeLa Kyoto cell lines. To generate RPE1 inducible cell lines recombination was initially selected for with 10 μg/mL puromycin for 72 hours followed by a stricter selection with 20 μg/mL puromycin for 72 hours prior to switching back to 10 μg/mL. The resulting KIF18A inducible cell lines were maintained in MEM Alpha (Life Technologies) with 10% FBS (Life Technologies) and 1 μg/mL puromycin (HeLa Kyoto) or 5 μg/mL puromycin (hTERT-RPE1) at 37°C, 5% CO_2_.

The resulting inducible cell lines were verified to be correct by extracting genomic DNA (QIAmp DNA Blood Mini Kit Qiagen #51106) and sequencing through the S284 mutation site (Eurofins). KIF18A inducible constructs were expressed with 2 μg/mL doxycycline (Thermo Fisher Scientific #BP26531).

### Immunofluorescence

Cells were seeded onto 12 mm glass coverslips in 24-well plates and fixed in -20°C methanol (Fisher Scientific, A412-1) or -20°C methanol with 1% paraformaldehyde (Electron Microscopy Sciences, 15710) for 10 minutes on ice. Coverslips were then washed three times for 5 minutes each in Tris-Buffered Saline (TBS; 150 mM NaCl, 50 mM Tris base, pH 7.4). Coverslips were blocked for 1 hour at room temperature in 20% goat serum in antibody dilution buffer (Abdil: TBS pH 7.4, 1% Bovine Serum Albumin (BSA), 0.1% Triton X-100, and 0.1% sodium azide). Coverslips were then washed two times in TBS for 5 minutes each prior to addition of primary antibodies. Primary antibodies were stored at -20 °C diluted 1:1 in glycerol and were subsequently diluted in Abdil at indicated concentrations and duration for staining. Coverslips were washed two times in TBS for 5 minutes each between primary and secondary antibody incubations. The following secondary antibodies conjugated to AlexaFluor 488, 594, or 647 were stored at -20 °C diluted 1:1 in glycerol and used at 1:500 dilution in Abdil for 1 hour at room temperature (Invitrogen Molecular Probes A11029, A11013, A11034, A11032, A11037, A11007, A11014, A11076, A21236, A21245, A21247). Coverslips were washed three times in TBS for 5 minutes each prior to mounting coverslips with Prolong Gold anti-fade mounting medium with DAPI (Invitrogen Molecular Probes, P36935).

### Microscopy

Fixed and live cell images were acquired on a Ti-E inverted microscope (Nikon Instruments) with a Clara cooled charge-coupled device (CCD) or Ti-2E inverted microscope (Andor) with a Prime Bsi sCMOS camera (Teledyne Photometrics) with a Spectra-X light engine (Lumencore). Both microscopes are driven by NIS Elements software (Nikon Instruments) and have an environmental chamber at 37°C. Images were acquired with the following Nikon objectives: Plan Apo 40x DIC M N2 0.95 NA, Plan Apo λ 60x 1.42 NA, or Apo TIRF 100x 1.49 NA. Images were processed and analyzed using ImageJ/Fiji (33,34).

### KIF18A Expression Level Quantification in HeLa Kyoto Inducible Cell Lines

After cells had been seeded onto 12 mm glass coverslips in 24-well plates for 24 hours endogenous KIF18A was depleted as described above and GFP-KIF18A was induced with 2 μg/mL doxycycline. 24 hours after knockdown and induction cells were fixed in -20°C methanol (Fisher Scientific, A412-1) for 3 minutes on ice and stained for rabbit KIF18A 1:100 (Bethyl, A301-080A), guinea Pig CENP-C 1:250 (MBL, PD030), and mouse gamma-tubulin 1:500 (Sigma Aldrich, T5326). All primary antibodies were incubated for 1 hour at room temperature, shaking. Twenty random mitotic cells were imaged for each condition per replicate, and expression level was quantified in ImageJ/Fiji by drawing an ROI around the mitotic spindle and measuring the KIF18A signal within that ROI. Background signal was determined by moving the same ROI to an area with no cells and measuring the GFP signal in that space. The mean background signal for each image was subsequently subtracted from the mean spindle KIF18A signal. All values were then normalized to the mean control background subtracted KIF18A intensity. Median and 95% confidence interval are reported for three individual biological replicates.

### Mitotic Index and Multipolarity Analyses

In a 24-well plate format, MDA-MB-231, HT-29, RPE1, or HeLa Kyoto cells were treated with control siRNA (5 pmol), KIF18A siRNA (5 pmol), DMSO, or Compound 3 (250 nM final concentration). After 24 hours, coverslips were fixed for staining using 1% paraformaldehyde (Electron Microscopy Sciences, 15710) in -20°C methanol. Cells treated with control siRNA, KIF18A siRNA, DMSO, or 250 nM Compound 3 were fixed and stained for mouse anti-γ-tubulin 1:500 (Sigma Aldrich, T5326), rabbit anti-KIF18A 1:100 (Bethyl, A301-080A), rat anti-α-tubulin 1:500 (Sigma Aldrich, MAB1864). All primary antibodies were incubated for 1 hour at room temperature. Twenty random fields of view were acquired for each condition and the number of mitotic cells as well as total cells were counted in each field of view. For multipolar counts, the number of poles was determined for each mitotic cell by comparing both the α-tubulin and γ-tubulin channels. Mean and standard deviations are reported from minimum three individual biological replicates for each condition.

### Chromosome Alignment and Spindle Length Analyses

Cells were seeded into 24-well plates and subsequently treated with control siRNA (5 pmol), KIF18A siRNA (5 pmol), DMSO, Compound 3 (250 nM) or Sovilnesib (250 nM). After 24 hours, coverslips were then fixed for immunofluorescence with 1% paraformaldehyde (Electron Microscopy Sciences, 15710) in -20°C methanol for 10 minutesCells were stained with mouse anti-γ-tubulin 1:500 (Sigma Aldrich, T5326), rabbit anti-KIF18A 1:100 (Bethyl, A301-080A), human anti-centromere antibody (ACA) 1:250 (Antibodies Inc., 15-235). All primary antibodies were incubated for 1 hour at room temperature except for the human ACA antibody, which was incubated at 4°C overnight. As described previously (23–25), single focal plane images with both spindle poles in focus were acquired. A boxed region of interest with a fixed height and width defined by the length of the spindle was used to measure the distribution of ACA-labeled kinetochore fluorescence using the Plot Profile command in ImageJ/Fiji. The ACA signal intensity was normalized internally to its highest value and plotted as a function of distance along the pole-to-pole axis. These plots were then fitted to a Gaussian curve and the Full-Width at Half Maximum (FWHM) for the Gaussian fit as well as the spindle length are reported for each cell analyzed. Mean and standard deviations are reported from minimum three individual biological replicates for each condition.

For assaying the effects of KIF18A mutants, inducible HeLa Kyoto cell lines were seeded into 24-well plates and subsequently treated with control siRNA (5 pmol) or KIF18A siRNA (5 pmol) and induced with 2 μg/mL doxycycline. After 24 hours, coverslips were then fixed for immunofluorescence with 1% paraformaldehyde (Electron Microscopy Sciences, 15710) in -20°C methanol for 10 minutes. Cells were then subsequently stained for rabbit GFP (Invitrogen, A11122) rat anti-α-tubulin 1:500 (Sigma Aldrich, MAB1864), and mouse anti-γ-tubulin 1:500 (Sigma Aldrich, T5326) for 1 hour at room temperature with shaking. As described previously (24,26), single focal plane images with both spindles poles in focus were acquired. A boxed region of interest with a fixed height and width defined by the length of the spindle was used to measure the distribution of DAPI labeled DNA fluorescence using the Plot Profile command in ImageJ/Fiji. The DAPI signal intensity was normalized internally to its highest value and plotted as a function of distance along the pole-to-pole axis using a custom macro in MatLab. These plots were then fitted to a Gaussian curve and the Full-Width at Half Maximum (FWHM) for the Gaussian fit, as well as the spindle length, are reported for each cell analyzed. Mean and standard deviations are reported from minimum three individual biological replicates for each condition.

### KIF18A Localization Analysis in Fixed Cells

RPE1, MDA-MB-231, and HT-29 cells were treated with either DMSO Compound 3 (250 nM), or Sovilnesib (250 nM) for 24 hours and then fixed for immunofluorescence. The following primary antibodies were used at the indicated dilutions: rat anti-α-tubulin 1:500 (Sigma Aldrich, MAB1864), rabbit anti-KIF18A 1:100 (Bethyl, A301-080A), and mouse anti-γ-tubulin 1:500 (Sigma Aldrich, T5326). Single focal plane images containing the γ-tubulin pole signal were acquired for individual cells. To measure KIF18A localization relative to the spindle pole, a 10-pixel wide line was manually drawn in Fiji from an individual γ-tubulin pole signal toward the center of the spindle. The profile intensities of KIF18A, α-tubulin, and γ-tubulin along that line was measured and recorded using the Plot Profile command in ImageJ/Fiji. Each of these profile intensities were normalized internally to its highest value. These normalized line scans were then aligned by peak γ-tubulin intensity and averaged for each pixel distance. Distance from pole to maximum KIF18A signal was determined for each aligned line scan. Mean and standard deviations are reported from a minimum of three independent experiments for each construct.

HeLa Kyoto cells for WT, S284A, and S284D cell lines were seeded onto coverslips and knocked down with lipofectamine RNAiMAX complexed with siRNAs as described above. Coverslips were simultaneously induced with 2 μg/mL doxycycline. 24 hours after knock down and induction, cells were fixed in 1% paraformaldehyde in ice-cold methanol for 10 minutes. The following primary antibodies were used at the indicated dilutions: rat anti-α-tubulin 1:500 (Sigma Aldrich, MAB1864), rabbit anti-GFP 1:1000 (Invitrogen, A11122), and mouse anti-γ-tubulin 1:500 (Sigma Aldrich, T5326). To measure KIF18A localization relative to the spindle pole, a 10-pixel wide and 6.5 μm long line was drawn in ImageJ/Fiji from an individual γ-tubulin pole signal toward the center of the spindle. The profile intensities of GFP, α-tubulin, γ-tubulin, and DAPI signal along that line were measured and recorded using the Plot Profile command in ImageJ/Fiji. The profile intensities of each were normalized to the highest value. Then, the normalized line scans were aligned to the highest γ-tubulin intensity and averaged for each micrometer distance. Lastly, the distances from the pole to the maximum KIF18A signal were calculated. A Kruskal-Wallis test with Dunn’s multiple comparisons test was performed.

### Proliferation Assays

MDA-MB-231, HT-29, or RPE1 cells were seeded into 24-well plates (Falcon, 353047) and allowed to adhere overnight before adding the indicated concentrations of Compound 3 or Sovilnesib. Approximately 6 hours after compound addition, plates were transferred to the Cytation 5 Cell Imaging Multi-Mode Reader (Biotek/Agilent) driven by Gen5 software (Biotek/Agilent). Plates were then imaged every 2 hours using a 4x Plan Fluorite 0.13 NA objective (Olympus) until confluent. Gen5 software was used to process images and measure cell counts at each time point using high-contrast brightfield imaging. Parameters used to determine cell count (cell size and light-intensity thresholds) were optimized for each cell line and previously validated (11). To determine the normalized fold change, the number of cells at the final timepoint was divided by the number of cells in the first time point and normalized to control treatment (DMSO) for each experiment. Data were generated from a minimum of three individual replicates. Proliferation Assays for HeLa Kyoto cell lines were performed in a similar manner. HeLa Kyoto cells inducibly expressing GFP, GFP-KIF18A wild type, GFP-KIF18A S284A, or GFP-KIF18A S284D were seeded into 24-well plates (Falcon, 353047) and allowed to adhere overnight. Knockdown of endogenous KIF18A was performed by adding 5 pmol KIF18A siRNA to each well. Simultaneously, expression of GFP or GFP-KIF18A constructs was induced by addition of 2 μg/mL doxycycline (Thermo Fisher Scientific, BP26531). To determine the normalized fold change, the number of cells at the final timepoint was divided by the number of cells in the first time point and normalized to GFP-KIF18A wild type for each experiment. Data were generated from minimum three individual replicates.

### Live Imaging of KIF18A Localization in RPE1 cells

RPE1 cells inducibly expressing siRNA resistant GFP-KIF18A were seeded into glass-bottom 24-well dishes (Cellvis, P24-1.5h-N). Cells were treated with 2 μg/mL doxycycline (Thermo Fisher Scientific, BP26531) to induce expression of GFP-KIF18A and 5 pmol KIF18A siRNA to deplete endogenous KIF18A. Approximately 1-1.5 hours before imaging, cells were switched into CO_2_-independent media (Gibco, 18045088) supplemented with 10% FBS and containing 100 nM SiR-Tubulin (Spirochrome, SC002), SPY595-DNA (1:10,000) (Spirochrome, SC301), 10 μM Verapamil (Spirochrome, SCV01) and 20 μM MG-132 (Selleck Chemicals, S2619). Images were acquired at 30 second intervals using a Nikon 60x 1.4 NA objective with 1 μm z-sections for 4-7 μm total (5-8 frames). To quantify changes in KIF18A fluorescence, a rectangular ROI encompassing the spindle was drawn and the KIF18A fluorescence within that ROI was measured and recorded using the Plot Profile command in ImageJ. The KIF18A fluorescence profile was measured for a timepoint before DMSO or Compound 3 addition (Initial: -1:30 minutes before DMSO or Compound 3 addition) and after DMSO or Compound 3 addition (Final: 12 minutes after DMSO or Compound 3 addition). Spindle Lengths were normalized on a 0 to 100% scale to account for differences in spindle lengths between cells. The ratio of Final to Initial KIF18A fluorescence was determined by dividing the KIF18A fluorescence values from the final timepoint by the initial timepoint for each individual cell. This ratio was then plotted as a function of % Spindle Length for each individual cell. To quantify the magnitude of the differences between the Final and Initial KIF18A fluorescence curves, the “Area Under Curve” calculation in GraphPad Prism was used.

### Microtubule-stimulated ATPase Assay

KIF18A motor activity was measured using an ADP-Glo luminescence assay (Promega, V9101). Pig microtubules (60μg/ml, Cytoskeleton, MT002) and ATP (25μM) were incubated with a 4-fold serially diluted test compound or DMSO in reaction buffer (15 mM Tris, pH 7.5, 10 mM MgCl2, 0.01% Tween20, 1% DMSO, 1 μM paclitaxel) at RT. Human KIF18A (1−417) protein (2.5 nM) was added to initiate the enzymatic reaction, the reaction mixture was incubated at RT for 120 min, and ADP-Glo reagents were added according to the manufacturer’s protocol. The luminescence intensity was measured using an EnVision plate reader (PerkinElmer). Percent inhibition for compounds = 100*(Positive control signal - sample signal) / (Positive control signal – Negative control signal), Curve fitting and IC_50_ values were determined by a 4-parameter nonlinear regression equation (variable slope) using GraphPad Prism 8.0 (GraphPad Software).

## Data Availability Statement

Data were generated by the authors and available on request

## ACKNOWLEDGMENTS

This work was supported by NIH R01GM130556 and NIH R35 GM144133 to JS, an NSF GRF 1842491 to KAQ, and a University of Vermont Summer Undergraduate Research Fellowship to OB.

---

**Figure S1:**
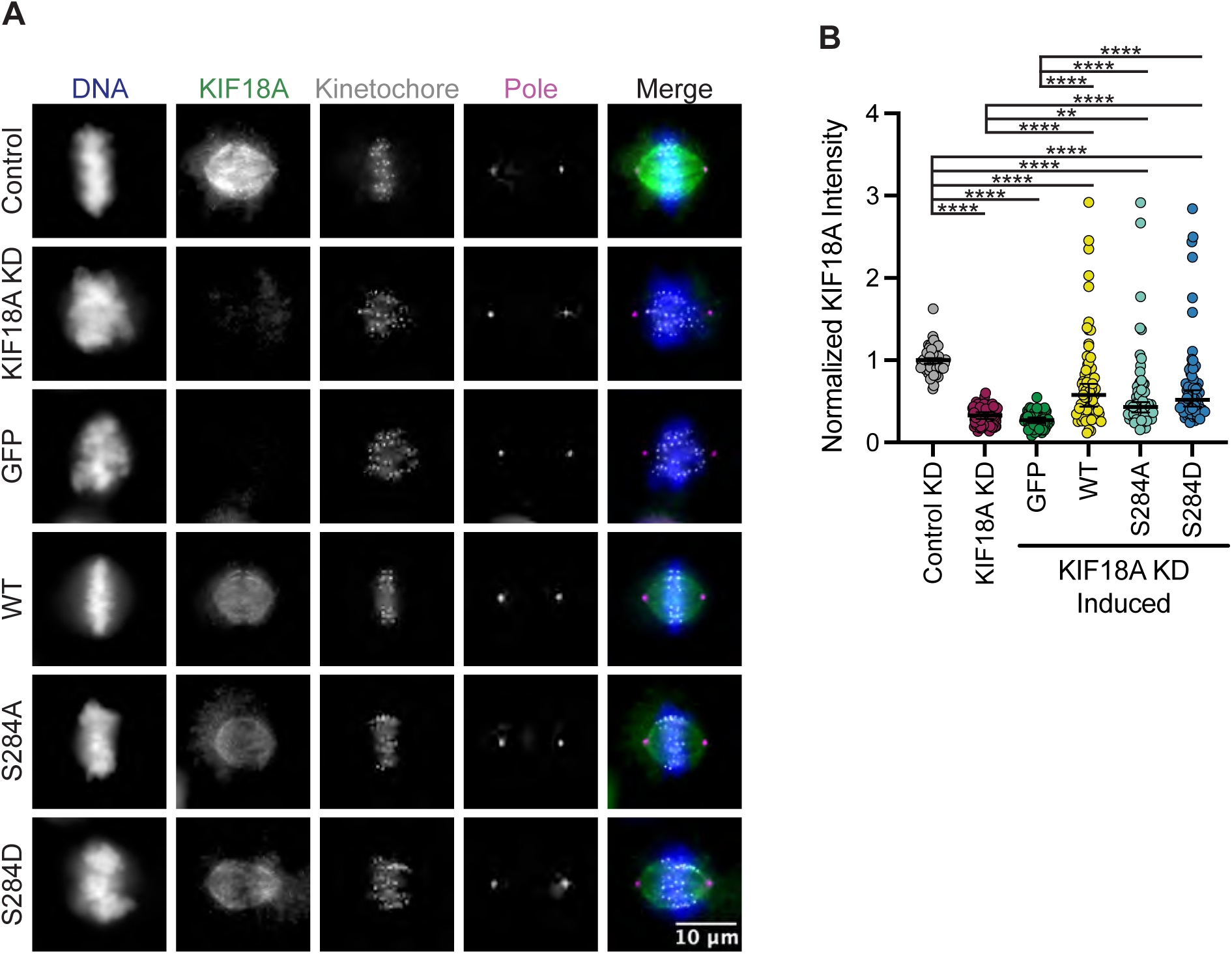
GFP-KIF18A S284 mutants are expressed at similar levels to wild type GFP-KIF18A. **(A)** Immunofluorescence images of HeLa Kyoto inducible cells fixed and stained 24 hours after endogenous KIF18A knockdown and induction of indicated GFP-KIF18A construct. Scale bar = 10 [)m. Text colors indicate pseudo color in merged image. From left to right DAPI/DNA, KIF18A antibody staining, CENP-C anti-body staining, α-Tubulin antibody staining, merged image. **(B)** Quantification of KIF18A expression from KIF18A antibody staining. Fluorescence values were to the mean control KIF18A intensity. Solid horizontal line indicates mean, vertical lines indicate standard deviation. Each dot represents a single cell. The total number of cells analyzed for each condition were control = 62, KIF18A KD = 80, GFP = 75, GFP-KIF18A WT = 68, GFP-S284A = 68, and GFP-S284D = 75. Data acquired from three experimental replicates. A one-way ANOVA with Tukey’s test for multiple comparisons was run, P values: < 0.05 (*), < 0.01 (**), < 0.001 (***), <0.0001 (****). If no significance is indicated, result was not significant (> 0.05).

**Figure S2:**
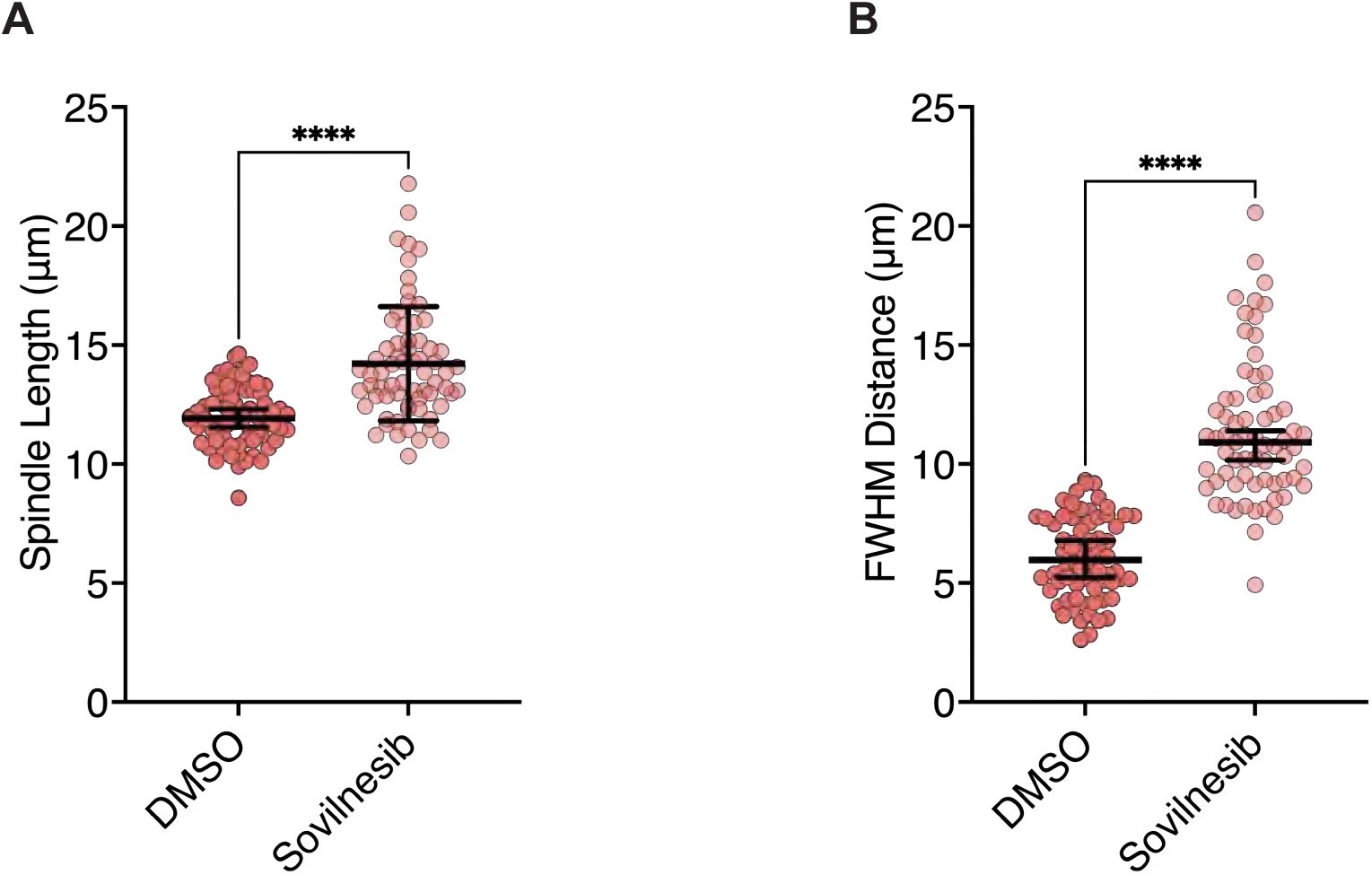
Sovilnesib mimics KIF18A knockdown phenotypes in MDA-MB-231 cells. Graphs of spindle lengths **(A)** and full-width at half maximum (FWHM) of centromere fluorescence distribution along the length of the spindle **(B)** measured in cells fixed and stained 24 hours after the indicated treatments. Each dot represents a single cell (N= 68 for DMSO, N=67 for Sovilnesib). Mean +/- standard deviation is displayed. Statistical results are shown for a Kruskal-Wallis with Dunn’s Multiple Comparisons test. P value style: < 0.05 (*), < 0.01 (**), < 0.001 (***), <0.0001 (****). n.s. indicates not significant (> 0.05).

**Figure S3:**
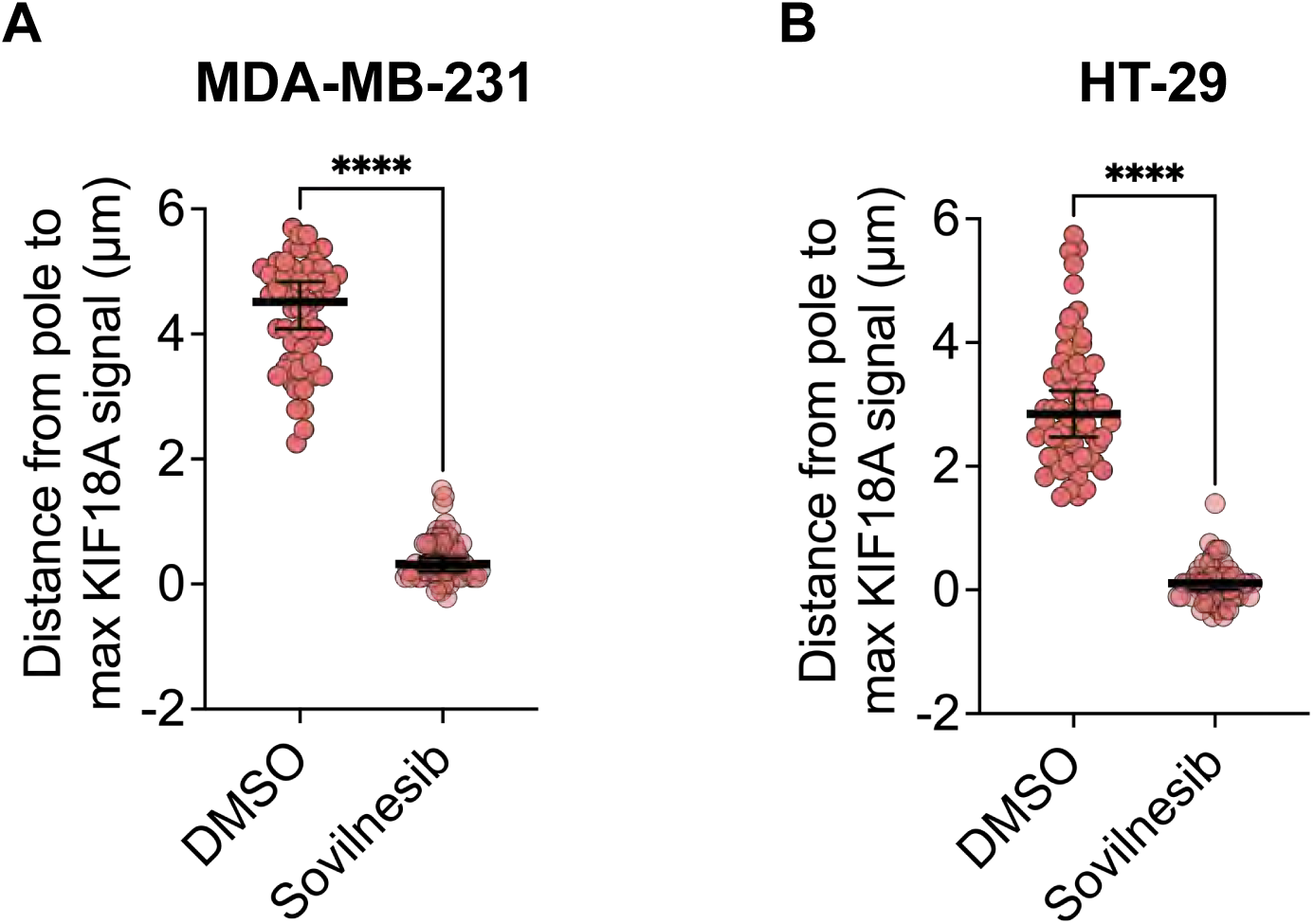
Sovilnesib disrupts microtubule plus-end localization of KIF18A. **(A-B)** Plots of distance from max KIF18A signal to the spindle pole derived from line scan analyses of KIF18A distribution in MDA-MB-231 **(A)** and HT-29 **(B)** cells. Each dot represents a single cell (MDA-MB-231: N=57 DMSO, N=60 Sovilnesib; HT-29: N=60 DMSO, N=60 Sovilnesib). Mean and SD are indicated by bars. Statistical results are shown for a Kruskal-Wallis with Dunn’s Multiple Comparisons test. P value: < 0.05 (*), < 0.01 (**), < 0.001 (***), <0.0001 (****). KIF18A localization analyses in fixed cell experiments showed accumulation of the motor at the spindle poles in KIF18A inhibitor treated cells, similar to KIF18A-S284 mutants. To confirm these results in live cells and determine the kinetics of KIF18A relocalization from microtubule plus-ends to spindle poles, we generated an hTERTRPE1 cell line that inducibly expresses GFP-tagged KIF18A (hereafter, RPE1 GFP- KIF18A). We arrested RPE1 GFP-KIF18A cells in metaphase with MG-132, then added DMSO or Compound 3 to track changes in KIF18A localization. In DMSO treated cells, GFP-KIF18A remained at the plus-ends of microtubules throughout the course of imaging (Figure 7A). In contrast, we observed loss of KIF18A from the plus-ends of microtubules and accumulation of KIF18A at spindle poles within minutes of Compound 3 addition (Figure 7A). This relocalization occurred in a pattern where a gap in KIF18A protein localization at the center of the spindle was observed a few minutes after compound addition and the gap then expanded towards the poles over time. We quantified these changes in KIF18A localization by plotting the GFP-KIF18A fluorescence profile across the spindle in the “Initial” timepoint (Pre DMSO or Compound 3 addition) and “Final” (12 minutes after DMSO or Compound 3 addition) timepoint, then calculated the ratio of the fluorescence values at the Final versus Initial timepoints along the length of the spindle (Figure 7B). To quantify the magnitude of changes in KIF18A localization over the 12-minute time course, the area under the curve for each ratio plot was also determined (Figure 7C). These analyses showed loss of GFP-KIF18A from the middle of the spindle and accumulation at the poles in Compound 3 treated cells, whereas DMSO treated cells did not exhibit these changes (Figure 7B-C). Combined, the fixed and live-cell imaging data demonstrate that chemical inhibition of KIF18A with Compound 3 treatment leads to relocalization of the motor from microtubule plus-ends to spindle poles in diploid and chromosomally unstable cell lines, similar to the localization defects observed for KIF18A-S284 mutants.

**Figure S4:**
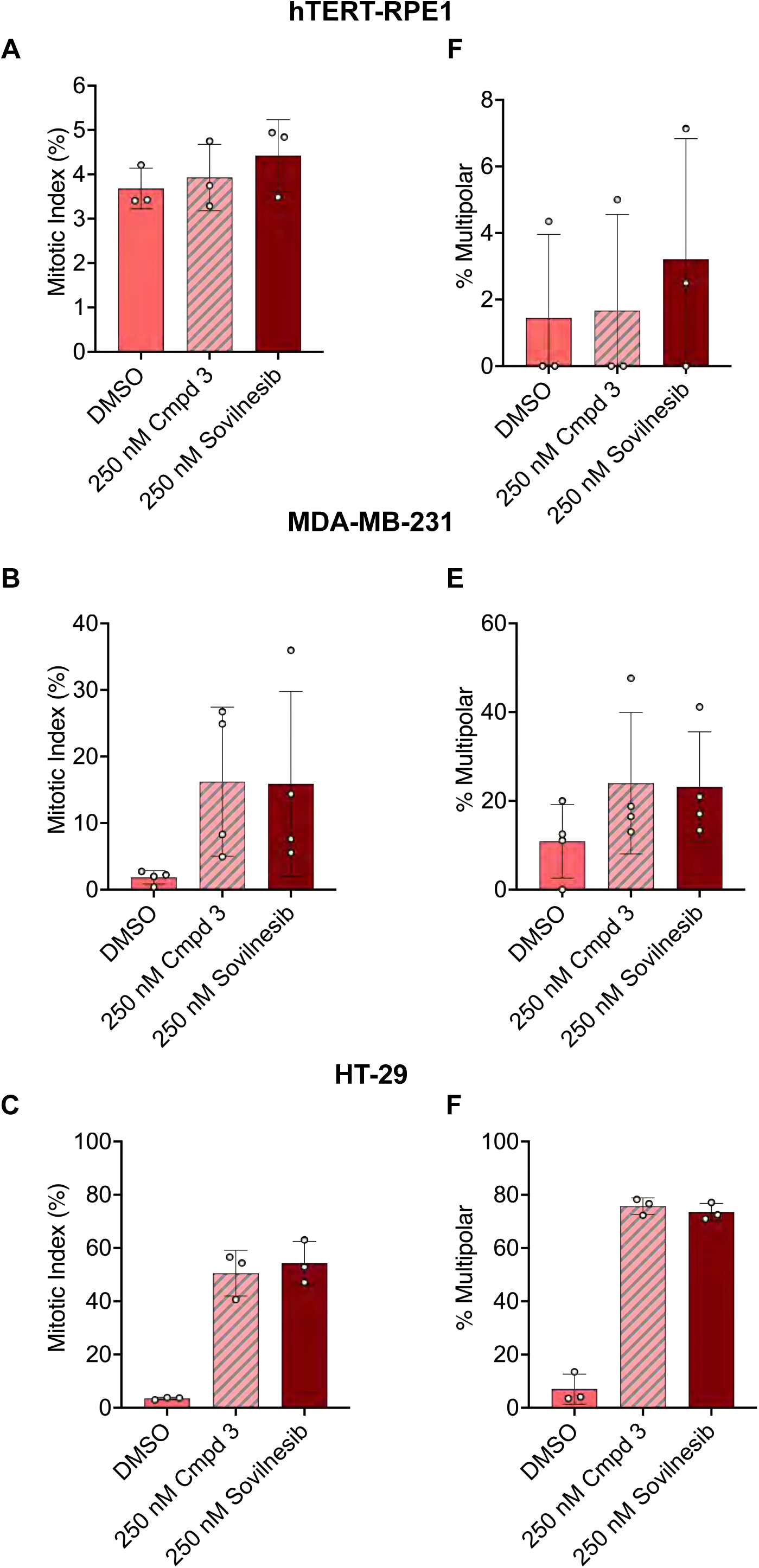
Sovilnesib treatment leads to mitotic arrest and multipolar spindles in chromosomally unstable breast and colorectal cancer cells. **(A-C)** Quantification of mitotic index (% of total cells in mitosis) in hTERT-RPE1 **(A)**, MDA-MB-231 **(B)**, and HT-29 cells **(C)** 24-hours after indicated treatments. Bars are mean +/- standard deviation. Each dot indicates an experimental replicate and data are from 3 independent experiments. **(D-F)** Quantification of multipolar spindles (% of total spindles) in hTERT-RPE1 **(D)**, MDA-MB-231, **(E)** and HT-29 cells **(F)** 24 hours after indicated treatments. Bars are mean +/- standard deviation. Each dot indicates an experimental replicate and data are from 3 independent experiments.

**Figure S5:**
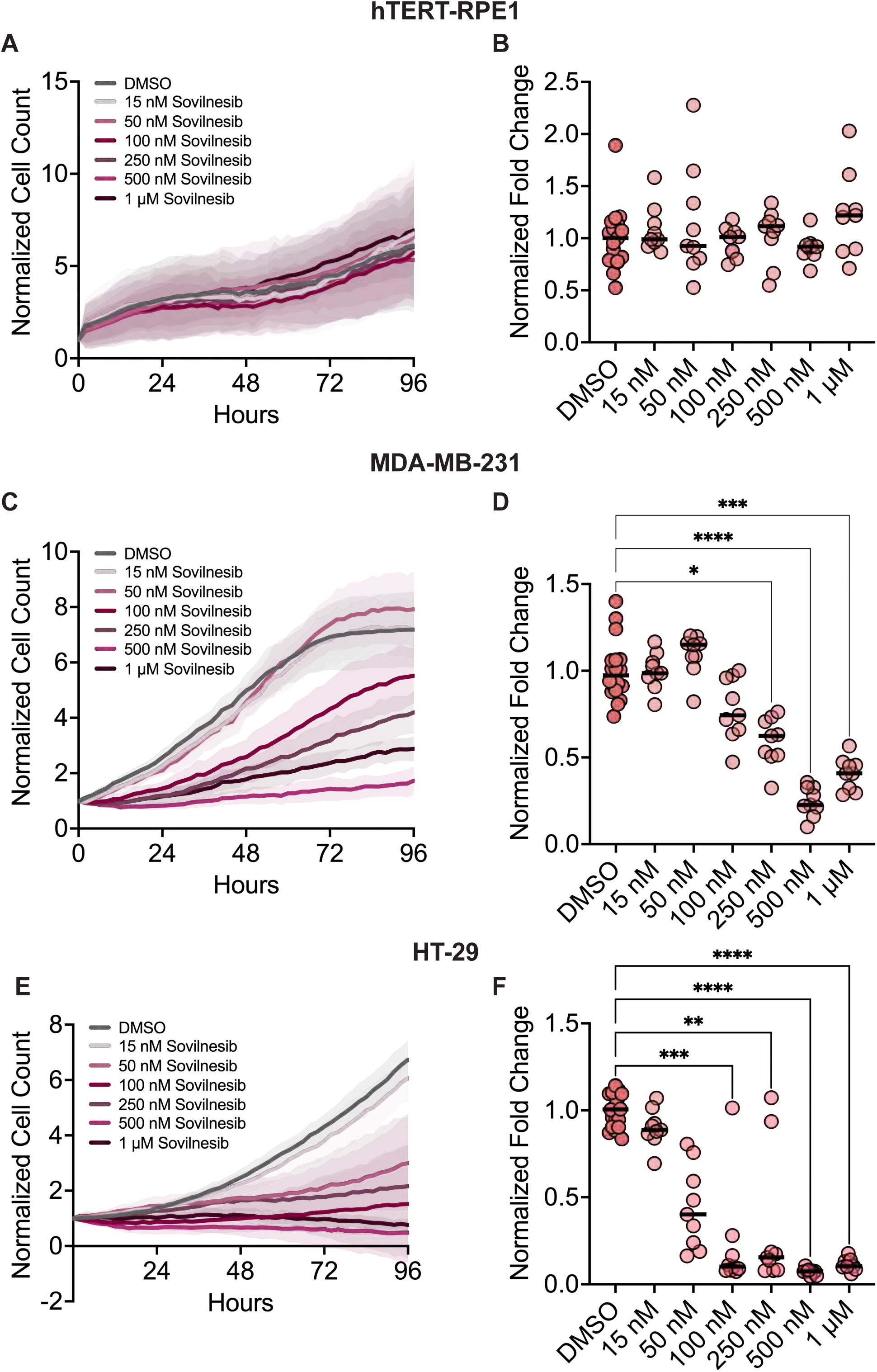
Sovilnesib reduces the proliferation of chromosomally unstable cells in a dose-dependent manner. **(A)** Plot of normalized cell count over time for cells treated with the indicated concentrations of Sovilnesib. Lines indicate mean and shaded area denotes SD. **(B)** Plot of normalized hTERT-RPE1 cell count (displayed as a % of DMSO control) as a function of Sovilnesib concentration. **(C)** Plot of normalized cell count over time for MDA-MB-231 cells treated with the indicated concentrations of Sovilnesib. Lines indicate mean and shaded area denotes SD. **(D)** Plot of normalized MDA-MB-231 cell count (displayed as a % of DMSO control) as a function of Sovilnesib concentration. **(E)** Plot of normalized cell count over time for HT-29 cells treated with the indicated concentrations of Sovilnesib. Lines indicate mean and shaded area denotes SD. **(F)** Plot of normalized HT-29 cell count (displayed as a % of DMSO control) as a function of Sovilnesib concentration. For plots in **(B)**, **(D)**, and **(F)**, each dot indicates an individual well and bars indicate mean values. Statistical results displayed from a Mann-Whitney test. P value style: < 0.05 (*), < 0.01 (**), < 0.001 (***), <0.0001 (****). n.s. indicates not significant (> 0.05). Data shown on all plots are from three independent experiments.

